# Genetic code expansion reveals site-specific lactylation in living cells reshapes protein function

**DOI:** 10.1101/2024.09.14.613019

**Authors:** Chang Shao, Shuo Tang, Siqin Yu, Chenguang Liu, Tianyan Wan, Zimeng He, Qi Yuan, Yueyang Zhang, Mengru Zhan, Hanqing Zhang, Ning Wan, Shihan Wu, Ren Xiang Tan, Haiping Hao, Hui Ye, Nanxi Wang

**Author notes:** These authors contributed equally to this manuscript (Chang Shao, Shuo Tang, Siqin Yu). Correspondence (H.Y.) (N.W.).

## Abstract

Still in its infancy, the functions of lactylation remain elusive. To address this, we established a comprehensive workflow for lactylation studies that integrates the discovery of lactylation sites with proteomics, the expression of site-specifically lactylated proteins in living cells via genetic code expansion (GCE), and the evaluation of the resulting biological consequences. Specifically, we developed a wet-and-dry-lab combined proteomics strategy, and identified highly conserved lactylation at ALDOA-K147. Driven by its potential biological significance, we site-specifically expressed this lactylated ALDOA in mammalian cells and interrogated the biological changes. We discovered that it not only inhibited enzyme activity but also elicited gain-of-function effects——it dramatically reshaped the functionality of ALDOA by improving stability, enhancing nuclear translocation and affecting gene expression. Further, we demonstrated broad applicability of this workflow to study distinct histone lactylation sites. Together, we anticipate its wide uses in elucidating causative links between site-specific lactylation and target-centric or cell-wide changes.

## Introduction

Protein lactylation is derived from lactate. It was first discovered on lysine residues of human histone proteins and was defined as an epigenetic mark that regulates gene expression in diverse biological contexts^1^. Recently, it was posited as a wide post-translational modification (PTM) that also modifies nonhistone proteins^2–7^. Exemplified by ALDOA, we found that ALDOA conservatively carries lactylation in its active site and therefore its activity is postulated to be inhibited by this PTM^3^. These representative studies have attracted enormous attention to the conceivably essential role of lysine lactylation during recent years and solicit researchers to seek answers for elusive questions such as: can lactylation instruct functional changes on the modified nonhistone proteins? Specifically, is lysine lactylation capable of inducing diverse biological consequences in living cells ranging from regulating modified protein’s activity, stability to fine-tuning interactions and even navigating proteins to certain compartments or organelles? Does lactylation installed at distinct lysine residues differentially control modified proteins?

However, to answer these questions of intense biological interests is highly challenging, as modification enzyme-based regulation is promiscuous and fails to site-specifically impose lactylation on the target protein in native biological contexts^8^. Furthermore, K to Q/E mutagenesis-based approaches^5, 7, 9, 10^ commonly used to mimic protein acetylation cannot fully mirror the lactylation state, nor can they distinguish between acetylation and lactylation of target proteins. This calls for technical advancements that allow precisely engineered lactylation. To address this demand, we and others have used GCE to site-specifically incorporate lactylated lysine (Klac) into model proteins, such as green fluorescent protein (GFP) and luciferase, in living cells with Klac-specific tRNA synthetase 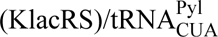 pairs^3, 11, 12^. Because this approach allows for convenient site-directed lactylation, we reason that it would be particularly suitable for inferring a causative link between lactylation at a specific site in a given protein of interest (POI) and its regulatory outcomes in a native and complex biological context (**Fig. 1a**). This allows better understanding of the biology of lactylation on nonhistone proteins, considering previous studies have mostly interrogated functionality of histone lactylation^1, 13–17^.

**Figure 1.**
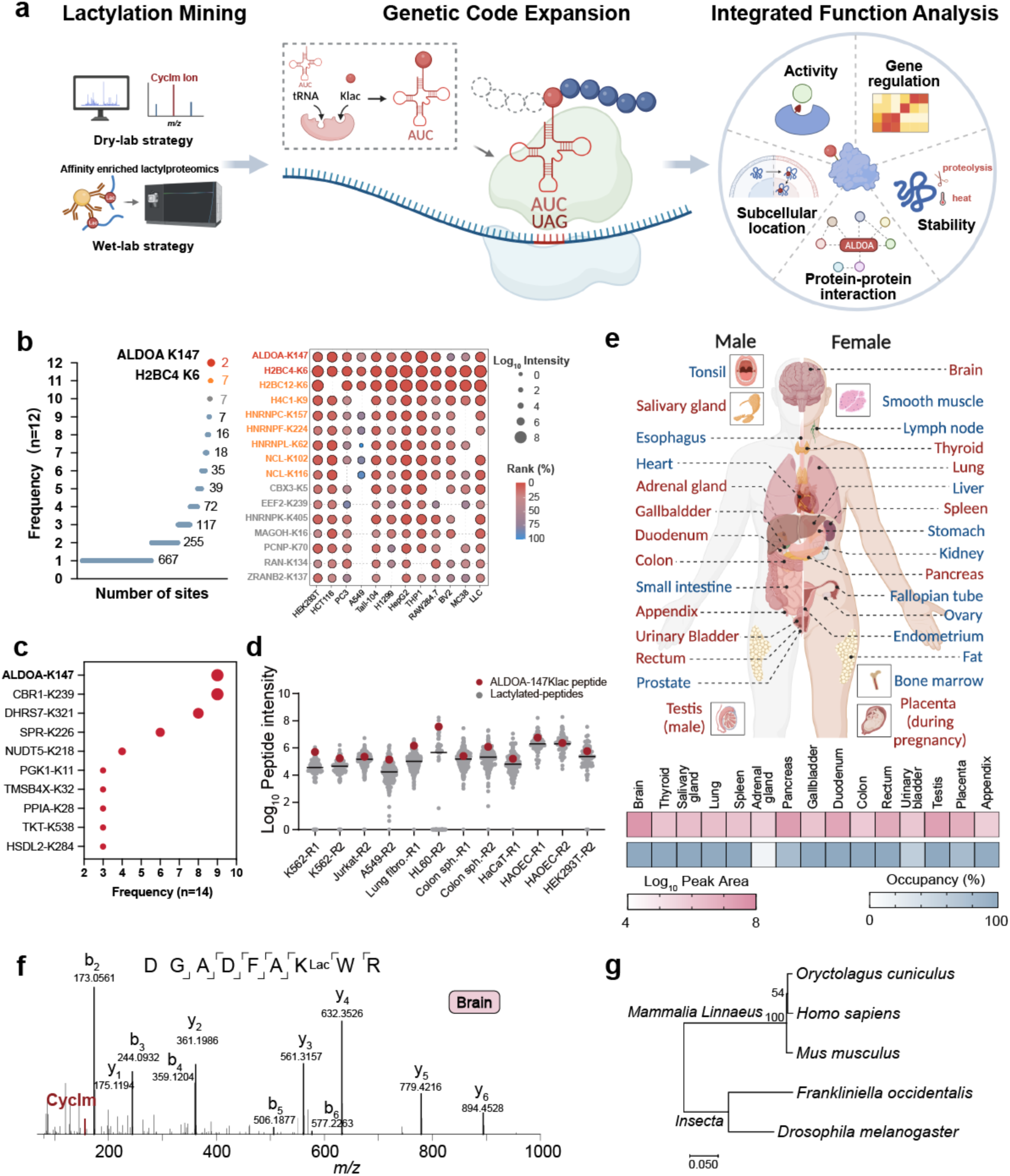
Proteomics mining pinpoints functionally important lactylation on ALDOA. (a) The workflow we have developed firstly uses proteomics to mine understudied lactylation sites of potential functional importance, and then uses GCE to express site-specifically lactylated proteins in living cells, followed by assembling a suite of biochemical tools to assess the biological consequences of site-specific lactylation. (b) Detection frequency of the lactylation sites in the affinity-enriched lactylproteome of representative human and mouse cell lines (left panel). Cells were treated with 25 mM lactate for 24 h to stimulate lactylation. The ion intensity (bubble size) and ranking (bubble color) of the identified lactylated peptides with a detection frequency of more than 10 times were shown in the corresponding bubble plot (right panel). (c) Detection frequency of the lactylation sites in the re-analyzed 14 human cell types retrieved from the Meltome Altas (PXD011929). (d) Abundance of the lactylated peptides (grey) in the re-analyzed human cell proteome retrieved from the Meltome Altas (PXD011929). Peptides carrying ALDOA-147Klac are highlighted in red. R1 and R2 represent the two biologically independent experiments we re-analyzed. (e) Abundance and occupancy of the peptides carrying lactylated ALDOA K147 in 15 healthy tissues (red) retrieved from a deep proteome atlas of 29 human healthy tissues (PXD010154). Occupancy was estimated based on the ratio of the abundance of the lactylated peptides divided by the sum of the lactylated and non-lactylated peptides at this site. (f) Representative MS/MS spectrum with the signature CycIm ion revealing lactylation on ALDOA-K147 in human brain (PXD010154). (g) Phylogenetic tree of amino acid sequences of ALDOA from the analyzed 5 species. The neighbor-joining method was used to generate the phylogenetic tree. Bootstrap values were calculated based on 1000 replications. Scale bar indicates the number of amino acid substitutions per site.

Here we first established a wet-to-dry-lab proteomics approach to mine functionally essential lactylation (**Fig. 1a**). We found that, besides affinity-enriched lactylproteome data repeatedly pinpointing lysine lactylation at residue K147 of ALDOA (ALDOA-147Klac), this modification was consistently identified in public proteome data of multiple human cell lines and tissue types using a cyclic immonium (CycIm) ion-based data mining strategy. Excitingly, ALDOA-147Klac also showed high occupancy in analyzed data and is even evolutionarily conserved in mammals and insects, suggesting its functional significance. With the developed GCE technique, we genetically engineered site-specifically lactylated ALDOA-K147Klac in HEK293T cells. Then, we established an integrated functional analyses (IFA) platform and collected a wealth of ALDOA-centric information, including enzyme activity, stability, subcellular location, gene regulation and interactions, to convey the effect of this modification on ALDOA (**Fig. 1a**). We found that single-site lactylation was sufficient to abolish the enzyme activity and inhibit the glycolytic flux. Surprisingly, we discovered that this single lactylation completely reshaped the function of ALDOA: it not only caused loss-of-function of ALDOA, but also enhanced the protein thermal stability, navigated ALDOA to the nucleus, and affected the transcriptome and its interacting partners. The workflow presented, by integrating powerful proteomics and GCE approaches with IFA (**Fig. 1a**), paves the way to establish causal links between the relatively nascent PTM—lactylation—and the biological consequences in living cells. Specifically, using ALDOA as an exemplary target, we demonstrate that lactylation on this metabolic enzyme reshaped its functions and modulated multiple tiers of biological processes expanding far beyond cellular metabolism.

## Results

### Proteomics mining pinpoints functionally important lactylation on ALDOA

Certain site-specific PTMs are prevalent in cells and serve as central players in core cellular processes^18, 19^. These are exemplified by PTMs such as histone acetylation/methylation that control transcription^20^ and phosphorylation on receptor tyrosine kinases that regulate cell growth and differentiation^21^. In contrast to these well-studied PTM states, lactylation has remained relatively understudied, and most functional studies have been devoted to histone lactylation^1, 13–17^. We therefore set out to discover additional nonhistone lactylation sites of biological significance.

We deployed a combined wet-and-dry-lab proteomics approach that first discovered affinity-enriched lactylation sites in a limited number of cell types (**wet-lab strategy**), followed by identifying the recurrent sites via mining large-scale public proteomic data resource collected from multiple types of human cells and tissues (**dry-lab strategy**). Using this combined strategy, we identified lactylated peptides from 8 human and 4 mouse cell lines and analyzed their frequency of occurrence among the cells tested (**Fig. 1b and Supplementary Table 1**). Among the identified lactylated peptides, ALDOA-147Klac was detected with the highest frequency, up to 100%. Furthermore, its intensities were consistently among the highest in almost all of the assayed cell lines (**Fig. 1b**). Naturally, the high prevalence and intensity of lactylated ALDOA-K147 implies its biological significance and intrigued us.

To confirm its conservative nature in the human proteome, we used our previously developed dry-lab experimentation strategy^3^ to screen for true lactylated peptides in unenriched human proteome data. The presence of the CycIm ion in MS/MS spectra of target-decoy database-searched lactylpeptides was set as the gold standard to signify true lactylation^3^. With this strategy, we reanalyzed lactylation sites from the public resource, Meltome Atlas (PXD011929)^3, 22^, and noted that K147 of ALDOA was once again the most commonly lactylated residue in the detected repertoire (**Fig. 1c**). We ranked the intensities of identified lactylated peptides and confirmed high abundances of the peptides bearing lactylated ALDOA-K147 (**Fig. 1d**). In agreement with the high ion intensities, lactylation on K147 is estimated to have an occupancy as high as 50.33% among the human cell lines examined^3^ (**Supplementary Fig. 1a**). In addition to human cell lines, we investigated the occurrence of site-specific lactylation in various human tissues. Excitingly, a deep proteome atlas of 29 healthy human tissues (PXD010154)^23^ revealed that ALDOA-147Klac was present in 15 of the analyzed tissue types (**Fig. 1e, Supplementary Fig. 1b**, **Table 2**). The MS/MS spectra retrieved for ALDOA-147Klac produced the CycIm ion, which is specific to intra-peptide lactylated lysine (**Fig. 1f**). Once again, this peptide exhibited both relatively high ion intensities and occupancy in human vital organs such as the brain, lungs and digestive system (**Fig. 1e and Supplementary Table 3**). Finally, we searched all publicly available affinity-enriched lactylproteome datasets together with unenriched proteomes of different species across the kingdoms of life^24^. We found that ALDOA-147Klac is conserved in human, mouse (our enriched proteomic data), rabbit (the kingdoms of life, PXD014877)^24^ and even insects, exemplified by fruit fly (Meltome Atlas, PXD011929)^22^ and western flower thrips (public lactylproteome, PXD030799)^25^ (**Supplementary Fig. 1c and Table 4**). We then constructed a phylogenetic tree based on the sequence alignment of ALDOA from these 5 species using MEGA X (**Fig. 1g**). We found ALDOA of *Homo sapiens* has a relatively close relationship with that of *Oryctolagus cuniculus* and *Mus musculus*, while showing a relatively distant relationship with those of the insects *Drosophila melaganoster* and *Frankliniella occidentalis*. As high levels of conservation are often indicative of important functions, this conserved lactylation across different species has aroused our keen interest. Together, the prevalence and high site stoichiometry of ALDOA-147Klac in humans, as well as its conservativeness across animal species, suggest that this site-specific lactylation is a functional hostpot and prompts further investigation into its biological consequences.

### Introducing site-specific lactylation in living cells with genetic code expansion

Intrigued by the biological changes caused by ALDOA-147Klac, we proposed to use GCE to produce site-specifically lactylated ALDOA in living cells and pursue the answer. Previously, we have evolved a pyrrolysyl (Pyl)-tRNA synthetase (PylRS) that specifically recognizes Klac (**Supplementary Fig. 2a**), which we named KlacRS1^3^, and managed to genetically encode lactylated target proteins and harvest the recombinant proteins in *E. coli*. In this work, we set out to engineer KlacRS with higher incorporation efficiency and stringency in order to introduce lactylation site-specifically into ALDOA in mammalian cells and investigate the biological consequences.

To achieve this goal, we first conducted positive selection with a focused *Methanosarcina mazei* PylRS (*Mm*PylRS) mutant library by completely randomizing residues Y306 and C348 to NNK (N=A/T/G or C, K=T or G). A hit with the Y306M and C348T mutations, exhibiting a Klac-dependent phenotype, was selected and named KlacRS2 (**Fig. 2a**). Meanwhile, inspired by the outstanding performance of chimeric tRNA synthetase/chimeric 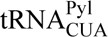 systems^26^, we created two chimeric KlacRS mutants, namely chKlacRS-WT (wild type) and chKlacRS-IPYE (IPYE mutant) by fusing the 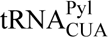 binding domain of *Methanosarcina barkeri* pyrrolysyl-tRNA synthetase (*Mb*PylRS-TD) or its active mutated form with the KlacRS1 catalytic domain at the C-terminus (KlacRS1-CD), respectively (**Fig. 2a**). To assess the amber suppression ability of KlacRS mutants described above, we co-expressed them with an enhanced GFP (EGFP) reporter carrying the Y39TAG mutation (EGFP-39TAG) in the presence or absence of Klac in *E. coli* cells. Interestingly, the results showed that in *E. coli* cells, although the orthogonality of KlacRS2, chKlacRS-WT, and chKlacRS-IPYE was enhanced compared with KlacRS1, KlacRS1 exhibited the highest Klac incorporation efficiency, with an expression yield reaching 39% of the WT EGFP (**Fig. 2b**).

**Figure 2.**
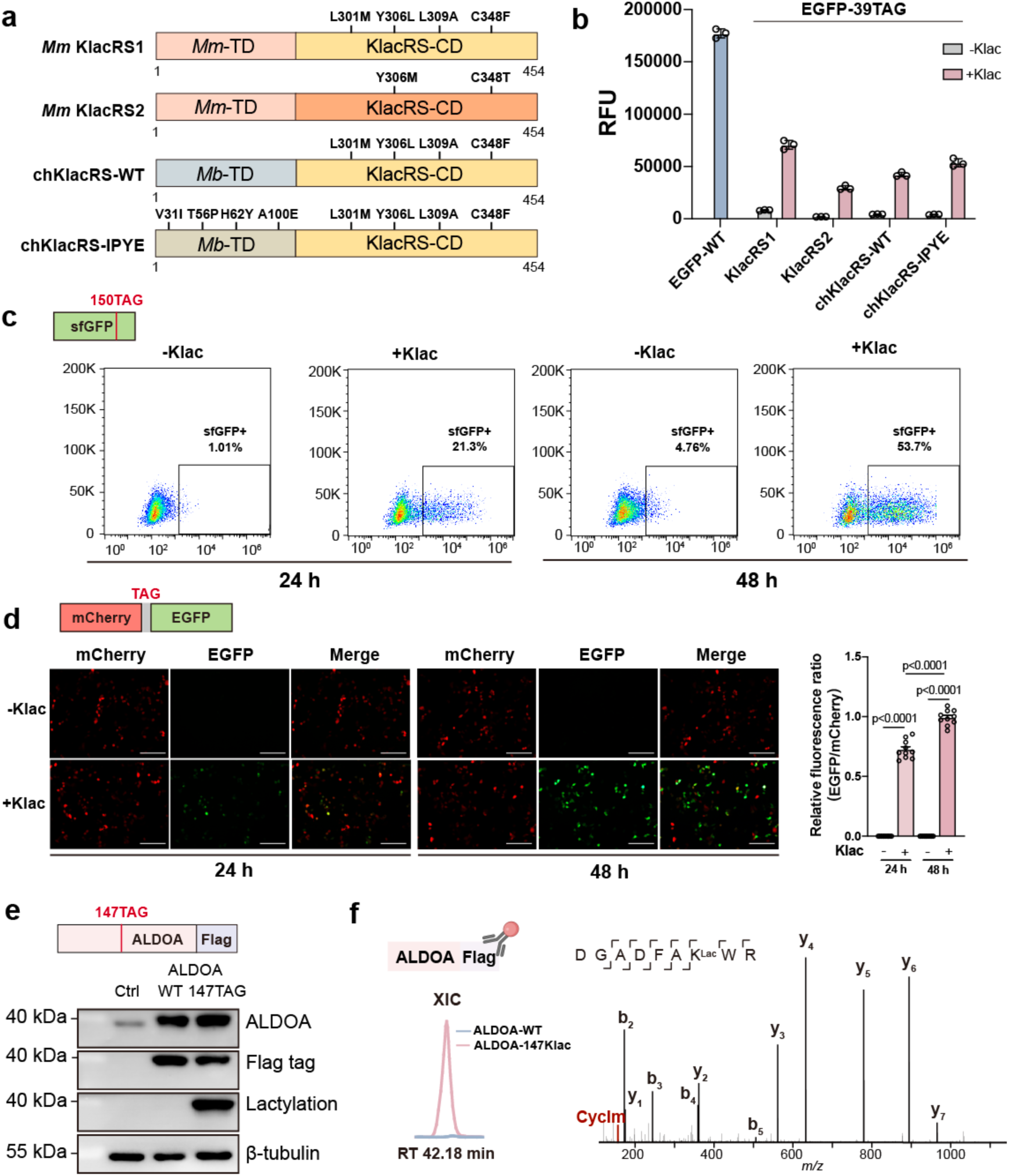
Introducing site-specific lactylation in living cells with genetic code expansion. (a) Cartoon structures of KlacRS variants for specifically encoding Klac. (b) Analysis of the incorporation efficiency of KlacRS variants by EGFP fluorescence assay. *E. coli* cells co-transformed with 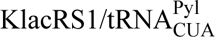 pair and EGFP-39TAG plasmids with or without Klac (1 mM, 16 h). Data represent the mean ± S.D. (n=3 biological replicates/group). (c) Flow cytometry analysis of Klac incorporation efficiency in HEK293T cells co-transfected with 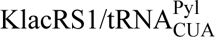 pair and sfGFP-150TAG plasmids treated with or without Klac (1 mM) for 24 h and 48 h. (d) Representative images of HEK293T cells co-transfected with KlacRS1 and 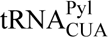 pair and mCherry-TAG-EGFP plasmids treated with or without Klac (1 mM) for 24 h and 48 h. The relative fluorescence ratio was calculated by comparing the mean intensity of EGFP to mCherry using ImageJ. Data represent the mean ± S.D. (n=10 biological replicates/group) and the p value was calculated by one-way ANOVA. (e) Immunoblotting analysis of HEK293T cells transfected with ALDOA-WT or co-transfected with 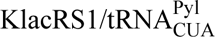 pair and ALDOA-147TAG plasmids treated with Klac (5 mM, 48 h). Abundance of overexpressed ALDOA was estimated using anti-ALDOA and anti-Flag antibodies. Genetically engineered ALDOA lactylation was detected using pan-lactylation antibody. (f) Validation of successful incorporation of lactylation on K147 of ALDOA. HEK293T cells transfected with ALDOA-WT or co-transfected with 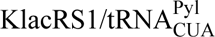 pair and ALDOA-147TAG treated with Klac (5 mM, 48 h), followed by immunoprecipitation using anti-Flag antibody and bottom-up proteomic analysis. Left, extracted ion chromatogram (XIC) of lactylated K147 peptide. Right, representative MS/MS spectrum with the signature CycIm ion revealing lactylation on ALDOA-K147.

Next, we examined the incorporation efficiency of KlacRS1 in mammalian cells. We co-transfected HEK293T cells with a plasmid expressing KlacRS1 and 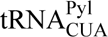 pair, and a plasmid containing sfGFP-150TAG reporter gene. Expression of full-length sfGFP was monitored by flow cytometry. As expected, strong sfGFP fluorescence was only detected when Klac was supplied, and the fluorescence intensity increased with longer incubation time (**Fig. 2c and Supplementary Fig. 2b**), suggesting that KlacRS1 can efficiently incorporate Klac in mammalian cells. To investigate the orthogonality of KlacRS1 in mammalian cells, we co-transfected a plasmid expressing KlacRS1 and 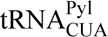 pair, and a dual-reporter plasmid encoding mCherry-TAG-EGFP gene into HEK293T cells. Expression of EGFP was monitored as a readout for amber suppression with or without Klac, while expression of mCherry was used as the transfection control. In the presence of Klac, strong mCherry and EGFP fluorescence were simultaneously observed by fluorescence microscope (**Fig. 2d**). In contrast, when Klac was omitted, no EGFP fluorescence signal was detected, indicating that KlacRS1 is highly active and orthogonal in mammalian cells. Notably, consistent with our previous observations, the relative fluorescence ratio of EGFP/mCherry increased by 27% when the incubation time was extended from 24 h to 48 h. Collectively, these results demonstrate that our KlacRS1 is capable of site-specifically incorporating Klac into target proteins in both *E. coli* and mammalian cells, and was therefore used for the following experiments.

Next, we optimized the Klac incorporation system in mammalian cells. Specifically, we measured EGFP fluorescence intensity of HEK293T cells co-transfected with KlacRS1 and 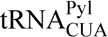 pair and EGFP-39TAG plasmids and treated with different concentrations of Klac. Both fluorescence imaging and flow cytometry revealed that full-length EGFP expression increased in a dose-dependent manner as Klac concentration was increased from 1 to 5 mM, reaching a plateau at 5 mM (**Supplementary Fig. 2c-d)**. Treatment with more than 5 mM Klac did not promote higher levels of EGFP expression (**Supplementary Fig. 2c-d)**, but resulted in notable cytotoxicity in HEK293T cells (**Supplementary Fig. 2e)**. Taken together, 5 mM Klac was selected for subsequent studies in order to achieve satisfactory incorporation efficiency while maintaining low cytotoxicity.

Using our optimized Klac incorporation system, we sought to site-specifically incorporate Klac into ALDOA in living cells via GCE. We co-transfected HEK293T cells with the 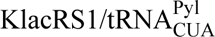 pair and ALDOA-147TAG plasmids. Immunoblotting analysis with the pan-lactylation antibody and Flag tag antibody confirmed successful expression of ALDOA-147Klac, and indicated that its expression level reached approximately 40% of ALDOA-WT (**Fig. 2e and Supplementary Fig. 3a**). Further, we immunoprecipitated both samples with anti-Flag antibody, followed by bottom-up analysis. Precise Klac incorporation at K147 of ALDOA was confirmed by detecting lactylated K147-bearing peptides only in cells genetically engineered to encode Flag-tagged ALDOA-147Klac, but not in cells encoding ALDOA-WT (**Fig. 2f**). Consistently, the non-lactylated K147-bearing peptide was only detected in the ALDOA-WT-encoding cells, but rather in ALDOA-147Klac-encoding cells (**Supplementary Fig. 3b**).

### Lactylation on ALDOA-K147 abolished enzyme activity and regulated glycolytic flux

Considering ALDOA is a metabolic enzyme responsible for the conversion of fructose 1,6-bisphosphate (FBP) into two triose phosphate, glyceraldehyde 3-phosphate (G3P) and dihydroxyacetone phosphate (DHAP), intuitively we asked whether lactylation would affect its activity due to the close proximity of the lactylated residue K147 to the C2 carbonyl group of FBP in the crystal structure of the enzyme/substrate complex^27^ (**Fig. 3a**).

**Figure 3.**
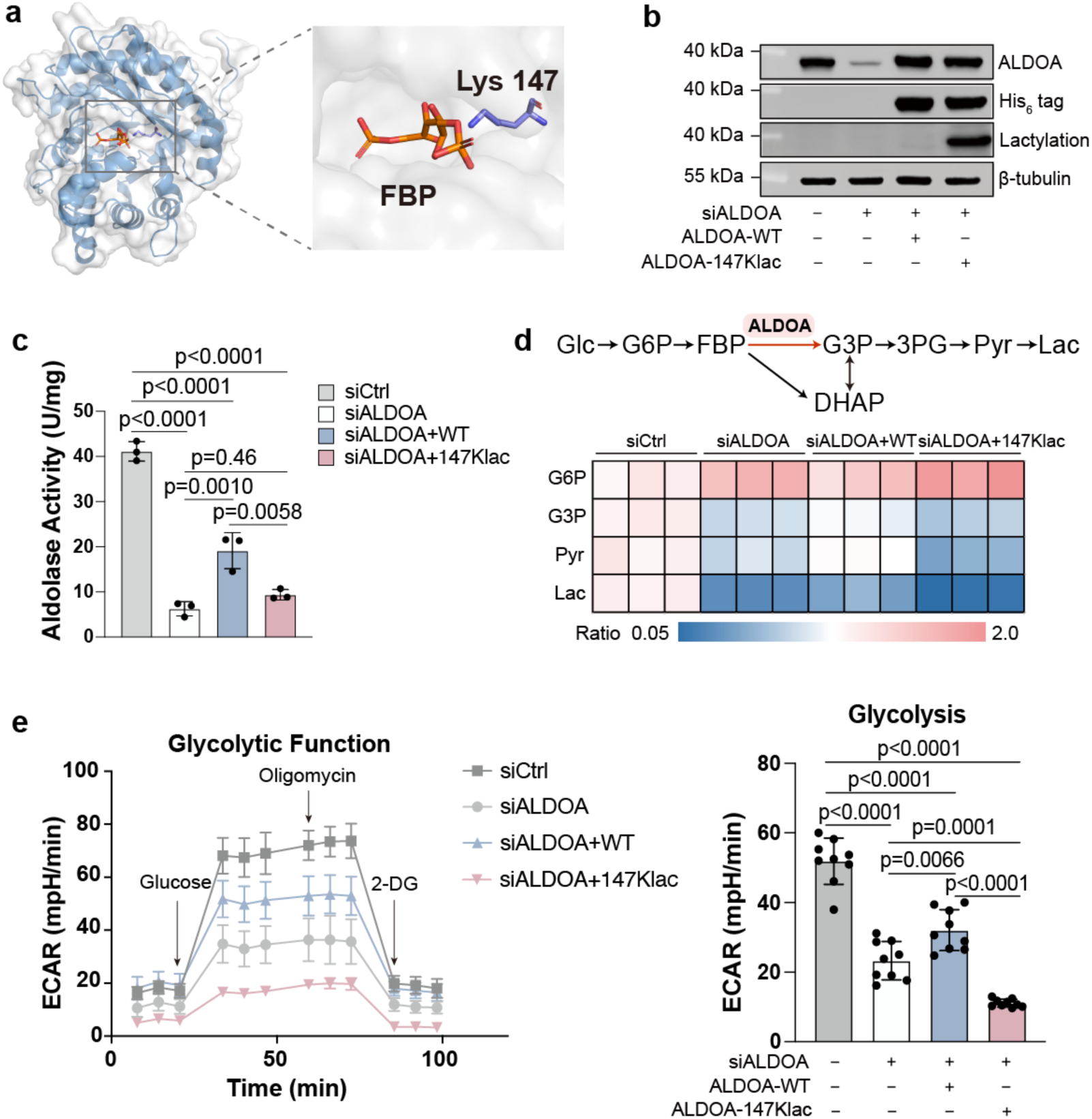
Lactylation on ALDOA-K147 abolished enzyme activity and regulated glycolytic flux. (a) Crystal structure of K147 in ALDOA (PDB 4ALD) and its substrate FBP. (b) Immunoblots of HEK293T cells expressing ALDOA-WT or ALDOA-147Klac after knocking down the endogenous ALDOA. (c) Aldolase activity of HEK293T cells expressing ALDOA-WT and ALDOA-147Klac after knocking down the endogenous ALDOA. Data represent the mean ± S.D. (n=3 biological replicates/group) and the p value was calculated by one-way ANOVA. (d) Heat map comparing abundance of metabolites involved in glycolysis. Abundance ratios were calculated by comparing ion intensities of individual metabolite in siALDOA, siALDOA+WT and siALDOA+147Klac groups, using the siCtrl group as a control. Abbreviations include: Glc, glucose; G6P, glucose 6-phosphate; FBP, fructose 1,6-bisphosphate; G3P, glycerol-3-phosphate; DHAP, dihydroxyacetone phosphate; 3PG, 3-phosphoglycerate; Pyr, pyruvate; Lac, lactate. (e) Seahorse analysis of HEK293T cells expressing ALDOA-WT or ALDOA-147Klac after knocking down the endogenous ALDOA. The ECAR was measured in real time under basal conditions and after the addition of glucose (10 mM), oligomycin (3 μM) and 2-DG (50 mM). Left, time course of a representative experiment. Right, determination of glycolysis rate. Data represent the mean ± S.D. (n=9 biological replicates/group) and the p value was calculated by one-way ANOVA.

Previously, we managed to site-specifically introduce lactylation into ALDOA in *E. coli*, and noted abolished enzyme activity with purified proteins^3^. Nevertheless, whether ALDOA-147Klac in living cells phenocopied this *in vitro* observation remains elusive. Here, as GCE enables producing ALDOA-147Klac in HEK293T cells, we are able to determine the effect of lactylation on ALDOA enzyme activity. We first knocked down the endogenous ALDOA via small interfering RNA (siRNA) to eliminate the interference from endogenous ALDOA (the siALDOA group), followed by overexpression of ALDOA-WT (the siALDOA+WT group) and the genetically encoded ALDOA-147Klac (the siALDOA+147Klac group), respectively (**Fig. 3b**). We found that ALDOA knockdown significantly impaired enzyme activity when comparing the siALDOA group with cells transfected with scramble siRNA (the siCtrl group). Subsequent analysis of the siALDOA+WT group and the siALDOA+147Klac group showed that the impaired activity can be partially restored by overexpression of ALDOA-WT; however, overexpression of ALDOA-147Klac cannot rescue the enzyme activity as ALDOA-WT (**Fig. 3c**). Considering that the abundance level of overexpressed ALDOA was similar between the two groups (**Supplementary Fig. 4a**), we confirmed that lactylation at K147 reduces ALDOA activity *in vivo*.

Given that ALDOA enzyme activity is vital for glycolysis, we hypothesized that dysfunctional ALDOA-147Klac may further interfere with glycolysis. We used metabolomics to evaluate the influence of lactylation on glycolysis by analyzing the siALDOA+WT and siALDOA+147Klac groups. We found that glucose 6-phosphate (G6P) — the glycolytic intermediate upstream of ALDOA—accumulated in the siALDOA+147Klac group compared to the siALDOA+WT group. G3P, pyruvate (Pyr) and lactate (Lac) — the metabolites downstream of ALDOA—were reduced in the former group compared to the latter group (**Fig. 3d**). Therefore, both the G3P/G6P and Lac/G6P ratios were reduced in the former group compared to the latter group (**Supplementary Fig. 4b**). The metabolomics data supported the disruption of glycolytic flux in living HEK293T cells due to site-specific lactylation on ALDOA-K147. To validate this, we compared the extracellular acidification rate (ECAR) between the ALDOA-WT and ALDOA-147Klac groups using Seahorse analysis. We confirmed that ALDOA knockdown decreased the ECAR level compared to normal cells. Again, ALDOA-WT overexpression partially rescued the decreased ECAR, while overexpression of ALDOA-147Klac failed to do so (**Fig. 3e**). These lines of evidences led us to conclude that site-specific lactylation at K147 inhibits the enzymatic activity of ALDOA, which results in disrupted glycolysis in living cells.

### Site-specific lactylation modulated ALDOA stability

Previous studies have demonstrated that PTM can not only alter enzymatic activity but also the biophysical properties of modified proteins^28^. One extensively studied property is protein thermal stability (PTS)^29, 30^. Changes in PTS could disrupt proteostasis and even cellular physiology^31^. The relationship between a specific PTM state and PTS has been inferred by comparing the melting curve of the modified proteoform with that of its unmodified counterpart^29^. Research has shown that site-specific phosphorylation on a serine residue located in the substrate binding domain of glyceraldehyde-3-phosphate dehydrogenase (GAPDH) can significantly destabilize it^29^, whereas in certain cases O-linked N-acetylglucosamine (O-GlcNAc) can stabilize the modified proteins^30^. These observations suggest that the lactylation of ALDOA may alter its thermostability. Therefore, we subjected the living cells encoding ALDOA-WT and ALDOA-147Klac to a temperature treatment following the Cellular Thermal Shift Assay (CETSA) workflow^32^. We noted that lactylation indeed displayed a thermostabilizing effect on ALDOA, as suggested by the melting curves of ALDOA obtained by immunoblotting against the Flag tag (**Supplementary Fig. 5a**).

As the resistance to proteolysis is another indicator of protein stability, we also assessed the proteolytic stability of ALDOA after lactylation using the drug affinity responsive target stability (DARTS)^33^ workflow. We treated cell lysates with a proteolytic enzyme and immunoblotted against the Flag-tagged ALDOA. We observed that the lactylated proteoform was significantly more resistant to proteolysis compared to unmodified ALDOA-WT (**Supplementary Fig. 5b**). This finding supports the lactylation-dependent stabilization detected by CETSA. Together, we anticipate that encoding lactylation in living cells via GCE may provide insight into the regulatory role of lactylation in protein stability and proteostasis, as well as exploring its implications in diseases^34^.

### Lactylation altered subcellular partition of ALDOA

Upon revealing lactylation on ALDOA-K147 as a loss-of-function PTM on enzyme activity, we sought to explore the gain-of-function capacities of this site-specific modification in regulating ALDOA. As PTM offers a common and dynamic method of modulating the subcellular localization of modified proteins, as exemplified by metabolic enzymes such as GAPDH^35, 36^ and pyruvate kinase M2 (PKM2)^9, 37^, we asked that whether the lactylation on ALDOA-K147 can alter the partitioning behavior of ALDOA. To test this hypothesis, we fused the ALDOA-WT and ALDOA-147TAG genes with an EGFP-His_6_ reporter at their C-terminus. This allowed for precise visualization and real-time tracking of ALDOA-WT and ALDOA-147Klac in living cells by confocal microscopy.

Initially, we discovered that ALDOA-WT exhibited a clear cytoplasmic distribution (**Fig. 4a**), which is consistent with its predicted localization via YLoc analysis^38^ (**Supplementary Fig. 6a**). This finding also agrees with previous reports of the primary cytoplasmic localization of ALDOA in A-431, U-251, and U2OS cells (retrieved from the Human Protein Atlas^39^, **Supplementary Fig. 6b**). Intriguingly, distinct from ALDOA-WT, fluorescent images showed that EGFP-tagged ALDOA-147Klac co-localized with Hoechst, indicating that ALDOA, after lactylation, is now distributed in the nucleus as well as the cytoplasm (**Fig. 4a**). Statistical analysis confirmed that lactylation promoted the subcellular translocation of ALDOA to the nucleus (**Fig. 4b**).

**Figure 4.**
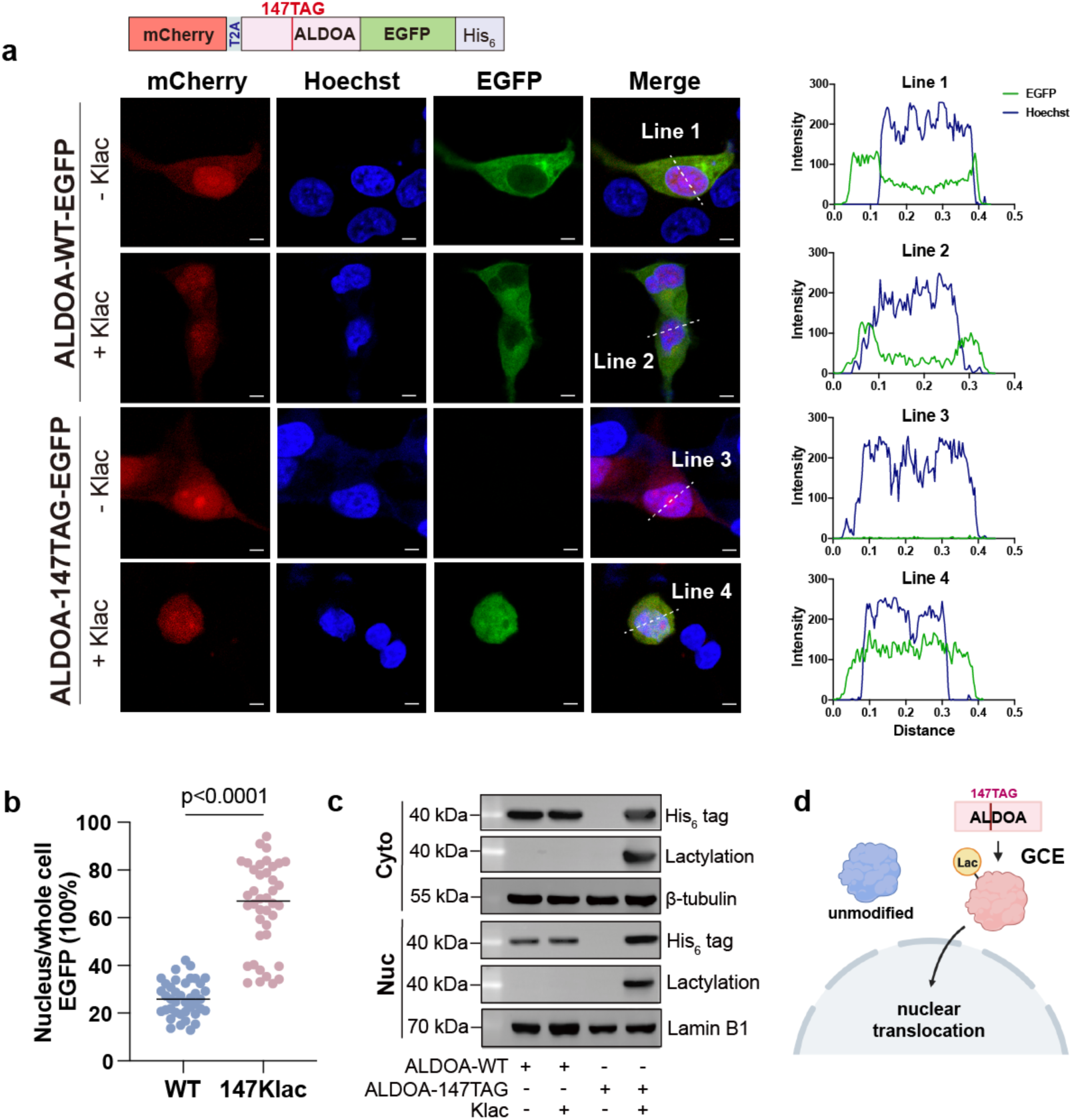
Lactylation altered subcellular partition of ALDOA. (a) Subcellular localization of EGFP-tagged ALDOA-WT (green) and EGFP-tagged ALDOA-147Klac (green) analyzed by confocal microscopy using cells expressing EGFP-tagged ALDOA-WT and ALDOA-147Klac, respectively. The nucleus was marked with Hoechst (blue). The mCherry (red) was used as the transfection control. Left, representative images showing ALDOA-147Klac partially translocated into the nucleus. Right, fluorescence intensity profiles across the indicated lines in the left. Scale bar, 50 μm. (b) Analysis of the mean fluorescence intensity of EGFP-tagged ALDOA in the nucleus compared to the whole cell using ImageJ for cells in a. n=40 biological replicates/group and the p value was calculated by Student’s t-test. (c) Subcellular localization of ALDOA-WT and ALDOA-147Klac determined by western blot analysis for cells co-transfected with 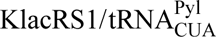 pair and ALDOA-147TAG plasmids or ALDOA-WT with and without Klac (5 mM, 48 h). Cytoplasmic β-tubulin and nuclear lamin B1 were used as controls. Cyto, cytoplasmic fraction; Nuc, nuclear fraction. (d) Schematic diagram showing that site-specific lactylation in ALDOA induced its subcellular translocation.

To confirm the nuclear accumulation of lactylated ALDOA observed through confocal microscopy, we conducted subcellular fractionation experiments. HEK293T cells were separated into cytoplasmic and nuclear fractions, and then were immunoblotted against His_6_-tag and lactylation. The results showed that ALDOA-WT primarily resided in the cytoplasm, while ALDOA-147Klac enhanced its translocation to the nucleus (**Fig. 4c**), consistent with the microscopic observations (**Fig. 4a**). Projecting forward, as lactylation can modify multiple lysines on the same protein and this might impart varied effects on protein subcellular localization, the GCE approach could be a valuable tool in resolving the relationship between different lactylation states and protein localization (**Fig. 4d**).

### Lactylation on ALDOA induced transcriptional changes in living cells

To uncover additional gain-of-function properties of lactylation on ALDOA, we reasoned that—as lactylation induced ALDOA to migrate into the nucleus—this might result in regulated gene expression based on previous findings on multiple metabolic enzymes exemplified by PKM2^37^, pyruvate decarboxylase (PDC)^40^ and methylenetetrahydrofolate dehydrogenase 1 (MTHFD1)^41^. Driven by this speculation, we performed RNA-sequencing (RNA-seq) analysis on HEK293T cells that site-specifically encode ALDOA-147Klac (147TAG+Klac) through co-transfection of 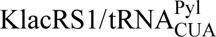 pair and ALDOA-147TAG plasmids and cultured with Klac, and used cells that were transfected in a similar manner but cultured without Klac as the control group (147TAG) (**Supplementary Fig. 7a**). Pairwise comparisons led us to identify 111 genes as differentially regulated (70 upregulated and 41 downregulated) by expression of ALDOA-147Klac (**Fig. 5a**). Gene ontology (GO) analysis indicates that these genes are associated with various Biological Processes (BPs) and Molecular Functions (MFs) (**Fig. 5b and Supplementary Fig 7b**). Specifically, most genes were enriched in the BP of regulation of cell junction assembly. In accordance with this, KEGG pathway enrichment analysis revealed the enrichment of genes in the cell adhesion pathway (**Supplementary Fig. 7c**). Furthermore, the analysis of Cellular Component (CC) showed that most of the regulated genes were enriched in the basolateral plasma membrane (**Supplementary Fig. 7d**). As these analyses appeared to pinpoint a plausible link between ALDOA-147Klac and cell adhesion/cell junction, we aimed to validate the hypothesis by focusing on the altered genes classified under the BP of cell junction assembly (**Fig. 5a-b**). We confirmed the transcript-level changes detected by RNA-seq with RT-qPCR analysis. This was exemplified by the down-regulation of pro-adhesion genes, including *CLDN1*^42^, *LRRTM2*^43, 44^ and *SLITRK6*^44^, and the upregulation of an anti-adhesion gene *GPBAR1*^45^ in cells encoding ALDOA-147Klac (**Fig. 5c and Supplementary Fig. 7e).**

**Figure 5.**
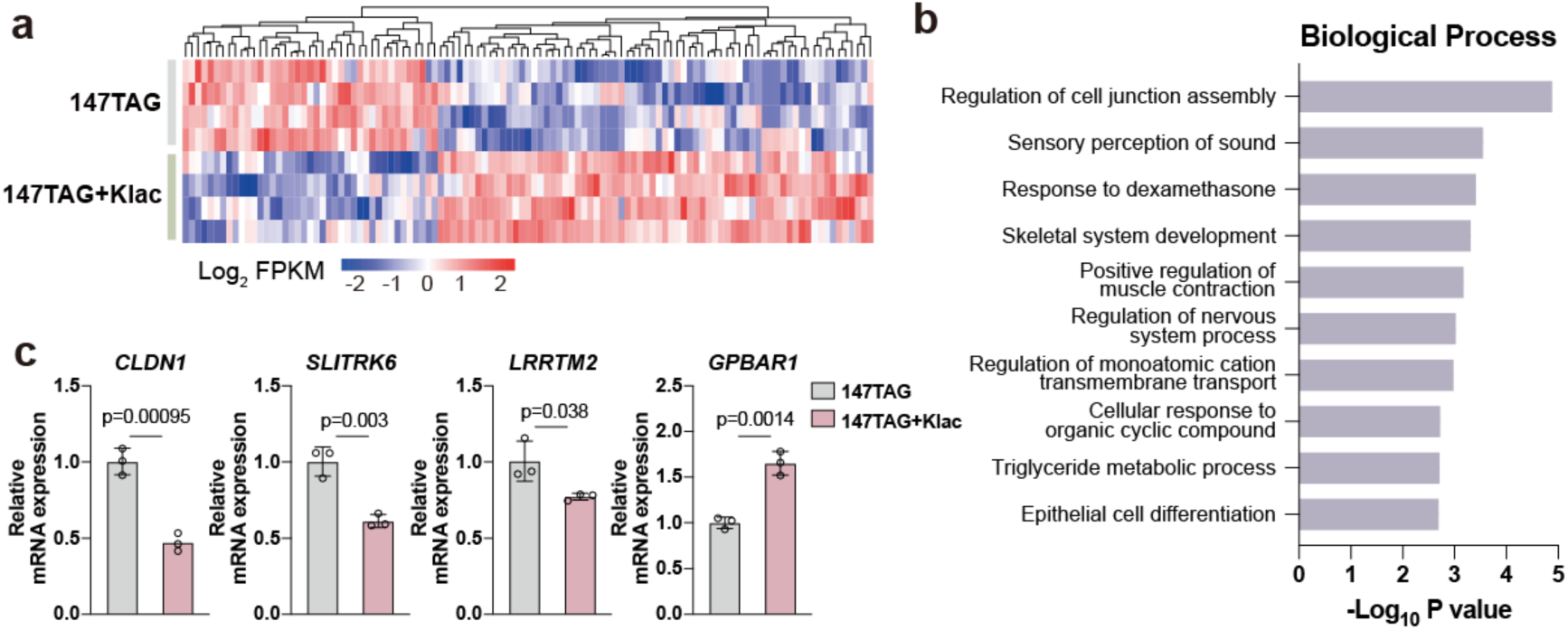
Lactylation on ALDOA induced transcriptional changes in living cells. (a) RNA-seq analysis of cells co-transfected with 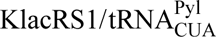 pair and ALDOA-147TAG plasmids with and without Klac (5 mM, 48 h). Heat map of genes showing significant changes (FC > 1.5 or < 0.67 and p value < 0.05 by Wald test as implemented in DESeq2) by plotting the log_2_FPKM value. FPKM, fragments per kilobase of transcript per million mapped fragments. (b) Bar plot of the GO BP analysis of differentially regulated genes in (a) from Metascape using the Benjamini-Hochberg correction algorithm. (c) RT-qPCR analysis of *CLDN1*, *SLITRK6*, *LRRTM2* and *GPBAR1* expression in cells co-transfected with 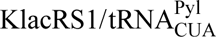 pair and ALDOA-147TAG plasmids with and without Klac (5 mM, 48 h). The endogenous β-tubulin gene was used as the internal control for normalizing target gene expression changes. Data represent the mean ± S.D. (n=3 biological replicates/group) and the p value was calculated by one-way ANOVA.

### ALDOA recruited different interacting partners after lactylation

Proteins often localize to different subcellular niches to fulfill their discrete functions. Therefore, as lactylation of K147 altered the subcellular localization of ALDOA, we speculate that this modification could also affect the interacting proteins recruited by ALDOA. To verify this, we performed co-immunoprecipitation (co-IP) experiments using ALDOA-WT- and ALDOA-147Klac-encoding HEK293T cells (**Fig. 6a**) and sought to identify the changed interacting proteins of Flag-tagged ALDOA after lactylation with quantitative proteomics. Quantitative assessment of the co-immunoprecipitated proteins revealed marked differences between the ALDOA-147Klac interactome and the ALDOA-WT interactome (**Fig. 6b and Supplementary Table 7**). GO analysis revealed that the interactomes of ALDOA-147Klac and ALDOA-WT are linked to distinct sets of BPs, MFs and KEGG pathways (**Supplementary Fig. 8a-c**). The analysis of CC further led us to note the enrichment of the ALDOA-147Klac interactome in the nucleus (**Fig. 6c**)—this agreed with the lactylation-promoted nuclear translocation of ALDOA (**Fig. 4**). Accordingly, the interacting proteins of ALDOA-147Klac were selectively enriched in certain BPs that primarily involve nuclear proteins. These BPs include regulation of cell cycle process, regulation of telomerase activity, mRNA transport and spindle organization (**Fig. 6d**). The interacting proteins of ALDOA-WT, which is primarily located in the cytoplasm, are enriched in metabolism-related BPs such as aerobic respiration and amide biosynthesis that also occur in the cytoplasm^46, 47^ (**Fig. 6d**). These results suggest that different proteins were recruited to ALDOA due to lactylation.

**Figure 6.**
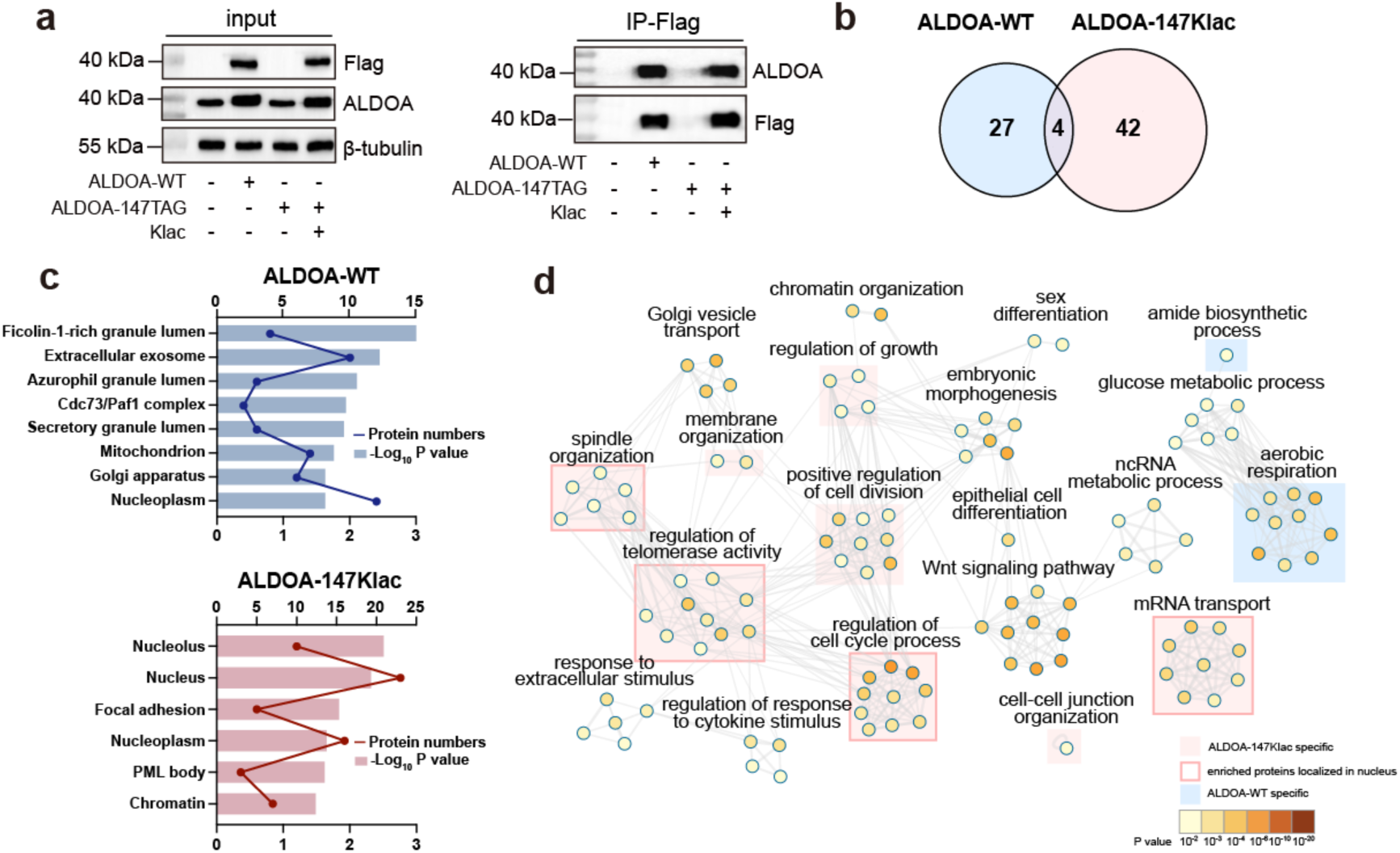
ALDOA recruited different interacting partners after lactylation. (a) Immunoblotting verified the enrichment of Flag tagged-ALDOA in ALDOA-WT and ALDOA-147Klac-expressed HEK293T cells. (b) Venn diagrams showing the interacting proteins of ALDOA-WT and ALDOA-147Klac. The interacting proteins of ALDOA-WT were enriched by co-IP using anti-Flag magnetic beads in HEK293T cells transfected with ALDOA-WT plasmid or vector, quantified by label-free quantification proteomics and calculated with a cut-off of FC > 2 and p value < 0.05 by Student’s t-test. The interacting proteins of ALDOA-147Klac in HEK293T cells co-transfected with 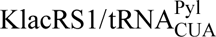 pair and ALDOA-147TAG with or without Klac (5 mM, 48 h) were enriched, quantified and calculated as the same procedure with ALDOA-WT. (c) GO CC analysis of the interacting proteins shown in (b) from DAVID bioinformatics website using the Benjamini-Hochberg correction algorithm. The lines indicate protein numbers enriched in each cellular component and the bars represent the −Log_10_ p value. (d) Network plot of significantly enriched GO BPs of interacting proteins of ALDOA-WT and ALDOA-147Klac shown in (b) using Metascape. Each node represents an enriched term and was colored by p value using the Benjamini-Hochberg correction algorithm. The node size is proportional to the number of input genes falling into that term. GO terms with a similarity > 0.3 were connected by edges and the thickness of the edge represented the similarity score. The network was visualized using Cytoscape. The red shade indicates that more than 70% of the proteins enriched in the specific BP were identified only in the ALDOA-147Klac group. If more than 70% of the enriched proteins in the red-shaded BPs are located in the nucleus, the shades are further framed in red. Blue shade is used to similarly indicate proteins specifically identified in the ALDOA-WT group.

### Site-specifically encoding histone lactylation in living cells

As we have demonstrated the capacity of GCE in genetically encoding lactylated ALDOA in living cells, we envisaged its expanded use in encoding site-specific histone lactylation, which is a well-demonstrated but still understudied epigenetic mark. Again, we first used proteomics to search for lactylation sites of potential biological significance by assessing their frequency of occurrence and ion intensities. Analysis of our detected lactylproteome of 8 human and 4 mouse cell lines revealed two lactylation sites in histone H4, namely lysine 9 (H4K9) and 13 (H4K13) with high ion intensity and frequency of occurrence compared to other histone sites (**Supplementary Fig 9a-b)**.

With these two target sites, we set out to use the GCE approach to generate histone H4-9Klac and H4-13Klac, respectively. After co-transfecting cells with plasmids encoding paired 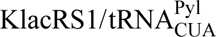 and H4-9TAG or H4-13TAG plasmids, respectively, we performed immunoblotting analysis using the anti-Flag antibody. Clear bands of full-length Histone H4 were observed in both groups of cells treated with Klac (+Klac) but not in groups without Klac (-Klac) (**Supplementary Fig 9c)**, indicating successful incorporation of Klac into Histone H4. Successful expression of Flag-tagged H4-9Klac and H4-13Klac in living cells was further confirmed through cross-validation using antibodies specifically recognizing H4-9Klac and H4-13Klac, respectively (**Supplementary Fig 9c)**. As such, we substantiated that the GCE approach can create cells capable of expressing histones with site-specific lactylation. This, along with epigenetic methods such as ChIP-Seq^1^ or cleavage under targets and tagmentation (CUT&Tag)^15, 48^, may represent a revolutionary tool for depicting the transcriptional consequences regulated by histone lactylation at single-residue resolution.

## Discussion

Lactate has recently been traced to the protein pool and found to be able to be derivatized into a nascent PTM, lactylation, on proteinaceous lysines^1^. This finding provides new insights into how cells sense changes in lactate levels and respond to this metabolic signal, potentially by adding lactylation to essential proteins. Projecting forward, ensuing functional interrogations of protein lactylation holds the promise of identifying markers or therapeutic targets for cellular malfunction and human diseases. However, current methods have limited this effort. In most cases, we pursue the biological consequences of a specific lactylation on its modified protein target by mutating the lactylatable lysine residue to a non-lactylatable residue^2, 4, 6^ and/or a lactylation-mimetic residue^5, 7^, followed by examining the resulting changes in protein function. Nevertheless, mutagenesis to natural amino acids cannot replicate the native structures and functions of lysine lactylation.

Alternatively, site-specific incorporation of unnatural amino acid (Uaa) in a particular POI via GCE allows for the expression of PTM-bearing proteins both *in vitro* and *in vivo*^49–55^. Such method has demonstrated its great value in deciphering various PTMs, such as acetylation^49^, phosphorylation^50^, sulfation^51, 52^, butyrylation^53^ and lipidation^54^. Unlike the PTMs described above, lactylation has rarely been genetically encoded into proteins other than model proteins and thus its potential remains largely untapped. We reasoned that as the number of available lactylome datasets continues to grow, the need for an efficient and precise GCE-based lactylation approach will increase. Herein, we presented the GCE approach for genetically encoding site-specific protein lactylation in living cells——this lays a foundation for further elucidating the regulatory roles of lactylation on POIs.

Considering that recent studies on lactylation’s functionality have mostly focused on histones and are therefore limited to transcriptional regulation^1, 13–17^, pursing the biological outcomes of nonhistone lactylation is anticipated to significantly enrich and expand our understanding towards lactylation. To decipher how lactylation regulates the function of its target protein and controls core biological processes *in vivo*, we created a comprehensive research workflow that integrates proteomic mining of lactylation sites and POIs, reconstitution of lactylated POIs in living cells via GCE, as well as IFA to assess site-specific lactylation-induced changes in POI’s activity, stability, interactions and cell-wide responses.

First, we developed a wet-and-dry-lab combined strategy to discover lactylation sites of biological importance through analyzing diverse proteome datasets. Our findings indicate that a particular lactylation, on ALDOA-K147, is present in all assayed human cell lines and most human tissues, as well as in lower non-human species such as rabbits and fruit flies. Given that the GCE approach allows producing site-specifically lactylated POIs, we used this strategy to hijack the native cellular translational machinery and successfully expressed the lactylated proteoform for the POI—ALDOA—in model cells, presenting initial efforts in unveiling lactylation functionality that expands beyond histones.

Using ALDOA as an exemplary POI, we first demonstrated that lactylated ALDOA-K147 inhibits the enzyme activity, thereby impairing glycolytic flux in living cells. This is consistent with our previous *in vitro* observation using purified ALDOA-147Klac, generated with the GCE method in *E. coli*^3^. Together, these lines of evidence demonstrate a causal relationship between lactate and glycolysis homeostasis——lactate can control over its biosynthetic pathway by inhibiting its upstream enzyme ALDOA through lactylation. This is not surprising as acylation installed in the active site of an enzyme often results in a loss-of-function in terms of enzymatic activity^56^.

Intriguingly, the IFA platform established in our workflow revealed that lactylation is also a gain-of-function modification. Specifically, we visualized that the site-specific lactylation on ALDOA-K147 altered subcellular partitioning of ALDOA and promoted its nuclear accumulation. Actually, in addition to lactylation, various PTMs such as acetylation^9^, succinylation^37^, phosphorylation^57^ and S-glutathionylation^35^ can also act as a relocalization signal for metabolic enzymes. However, the mechanisms underlying nucleocytoplasmic translocation following lactylation, as in the case of ALDOA, are not fully understood. The GCE approach adopted in this lactylation research workflow, by enabling the production of precisely lactylated POIs in cells, may facilitate the investigation of the dependent machinery. For instance, we can selectively silence or inhibit the components hypothetically involved in this process, such as importins and exportins^58^, and then monitor the subcellular partitioning of lactylated ALDOA. In addition to the transport machinery, it is also possible to elucidate the regulatory machinery of lactylation^1, 59^. The candidate “writer”/ “eraser” of POI can be conveniently identified by treating the lactylation-encoding cells with genetic or pharmacological intervention that selectively target the candidate “writer”/“eraser” enzyme and assessing the changes in lactylation levels.

Intuitively, we next determined transcript-level changes in cells expressing the nucleus-localized ALDOA-147Klac, as relocalization to the nucleus often enables proteins to participate in gene transcription^60, 61^. We noted affected transcription of certain genes involved in the BP of cell junction. This is particularly interesting as it supports previous findings that metabolic enzymes, typically localized in cytoplasm or mitochondria, can perform non-metabolic functions when redistributed to the nucleus^41, 57^. These “moonlighting” enzymes then govern a wide spectrum of instrumental cellular processes, including influencing histone PTM patterns, interacting with transcription factor, altering chromatin structure and mRNA stability—these diverse mechanisms converge in regulated gene transcription^61^. Thus, we propose that, by using the GCE approach, an accurate roadmap linking the subcellular localization of POIs, along with the dependent biological consequences such as gene transcription, to site-specific lactylation can be created in the future.

Besides incorporating Klac in understudied lactylated substrates like ALDOA, we demonstrated the applicability of the GCE approach in encoding lactylated histones. Given the essentiality of histone PTMs in controlling epigenetic phenomena, we envision that our approach allows attaining transcriptional outputs regulated by histone lactylation at single residue-resolution. Such knowledge, together with a wealth of information regarding changed gene expression in response to classic histone acylations such as acetylation^18^, is anticipated to explore the indexing potential of histone code and uncover the specific downstream outcomes that stem from recognition of distinct histone codes^62^.

Several limitations of the current GCE method of lactylation incorporation in living cells exist and warrant future optimization. First, our method using the orthogonal 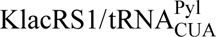 pair, although enabling precisely incorporating lactylation in living mammalian cells, is based primarily on transient transfection^11, 12, 51, 52, 54^. This limits the scope of genetically encoded lactylation only to cell lines that allow efficient transfection. Future adaptation of the GCE method with PiggyBac transgenesis^49^ or CRISPR-based genome editing^53^ is anticipated to generate stable cell lines with efficient and uniform incorporation of lysine lactylation. Second, we employed bottom-up proteomics to uncover the lactylation sites naturally present in cells and direct the appropriate sites for Klac incorporation. Nevertheless, bottom-up analysis is competent in detecting site-specific PTMs through proteome-wide analyses, but cannot measure intact proteoforms of POIs and hence suffers from the loss of PTM combinatorial information^63^. Continuous efforts will be made to use the complementary proteomics approaches to uncover how lactylation may be installed at distinct residues of POIs simultaneously, well-exemplified by histone H4 in this study, and work in combination to modulate protein functions. It is also possible that lactylation co-exist with other PTMs on the same POI^64^; the ability to reveal how these PTMs crosstalk and act in a concerted manner to regulate protein functionality in living cells will deepen our understanding of lactylation from a comprehensive view of the PTM landscape than studies focusing on lactylation alone. Therefore, this warrants future efforts towards developing GCE methods that enable simultaneous incorporation of lactylation and other PTMs in POIs.

In summary, we have developed a research workflow for lactylation that assembles proteomic mining, the GCE approach, and IFA. This workflow is amenable to study lactylation of any POIs in living cells. It equips us with the ability to accurately assess the perturbed activity, stability, spatial distribution, interacting proteins of POIs and cell-wide changes caused by site-specific lactylation. The developed workflow is expected to spur research in the field of lactylation by helping to resolve the enigmatic functionality of site-specific lactylation in cells and even organisms. Such knowledge will open up the possibility of investigating whether enhancing or inhibiting specific lactylation could be a therapeutic approach for diseases, particularly those associated with metabolic disorders.

## Acknowledgements

This research was supported by the National Natural Science Foundation of China (grant 82104050 and 22377058 to N.W., grant 82173783 to H.Y.), the Natural Science Foundation of Jiangsu Province (BK20220088, BK20210692), the National Key Research and Development Program of China (2021YFA1301300), the Fundamental Research Funds for the Central Universities (2632022YC03), the Overseas Expertise Introduction Project for Discipline Innovation (G20582017001).

## Author contributions

H.Y. and N.W. (Nanxi Wang) conceived the project. C.S., S.T., S.Y., R.X.T., H.H, H.Y. and N.W. (Nanxi Wang) designed the experiments. C.S., S.Y., Z.H. and N.W. (Ning Wan) performed the proteomics experiments and lactylation mining. S.T. contributed to plasmid construction. C.S., S.T. and Q.Y. performed the flow cytometry experiments. C.S., S.Y., S.T., C.L. and Y.Z. performed fluorescence and confocal imaging experiments. C.S. and S.Y. conducted metabolomics and seahorse experiments. Y.Z. performed DARTs and CETSA experiments. C.S. and Q.Y. contributed to RNA-seq and RT-qPCR analysis. C.L., Y.Z. and H.Z. contributed to co-IP analysis. S.T. and M.Z. contributed to KlacRS screening. T.W. and S.W contributed to Klac synthesis. H.Y., C.S. and N.W. (Nanxi Wang) wrote the manuscript.

## Materials & Correspondence

Correspondence and material requests should be addressed to (lead contact).

## Methods

### Cell culture and chemicals

THP-1 and MC38 were purchased from Cell Bank/Stem Cell Bank, Chinese Academy of Sciences (Shanghai, China). HEK293T, HCT116, PC3, H1299, HepG2, A549, TALL-104, RAW264.7, BV2 and LLC cells were purchased from American Type Culture Collection (ATCC) and cultured at 37 ℃ in a 5% CO_2_ atmosphere. HEK293T, HepG2, RAW264.7, BV2 and LLC were cultured in Dulbecco’s Modified Eagle Medium (DMEM). HCT116, PC3, A549, H1299, THP-1 and MC38 were cultured in RPMI-1640 medium. TALL-104 was cultured in RPMI-1640 medium containing 10 ng/mL interleukin-2 (PeproTech, cat. no. 200-02). All culture media were purchased from Gibco and supplemented with 10% fetal bovine serum (Excell, cat. no. FSP500), 100 U/mL penicillin and 1 μg/mL streptomycin (Thermo Fisher Scientific, cat. no. 15070-063). LC-MS grade water and acetonitrile (ACN) were obtained from Merck (Darmstadt, Germany). All chemicals were purchased from Sigma-Aldrich unless otherwise specified. The oligonucleotide primers were obtained from Tsingke (Nanjing, China).

### Dry-lab strategy for lactylation mining

For lactylation mining based on public proteome data, database searching parameters were set according to the mass spectrometer and parameters used in the literature. Specifically, Meltome Atlas data^22^ were searched using PEAKS Studio XPro (Bioinformatics Solutions Inc.) against the UniProt human proteome database (UniProt_Human_reviewed_29-11-2021.fasta). A mass tolerance of 5 ppm was allowed for precursor ions and 0.02 Da for fragment ions. Human tissues proteome data^23^ were searched using PEAKS Studio online 11 against the UniProt human proteome database (UniProt_Human_reviewed_20-7-2023.fasta). A mass tolerance of 10 ppm was allowed for precursor ions and 0.05 Da for fragment ions. Proteome data collected from different animal species were searched using PEAKS Studio Xpro against their species-specific UniProt FASTA database. Specifically, UniProt_Human_reviewed_15-5-2023.fasta for *Homo sapiens*, UniProt_Mouse_reviewed_11-6-2023.fasta for *Mus musculus*, UniProt_Rabbit_reviewed_26-6-2023.fasta for *Oryctolagus cuniculus*, UniProt_Fruit fly_reviewed_27-6-2023.fasta for *Drosophila melanogaster* and UniProt_Western flower thrips_unreviewed_7-12-2023.fasta for *Frankliniella occidentalis* were used. A mass tolerance of 7 ppm was allowed for precursor ions and 0.02 Da for fragment ions. For other parameters, we set trypsin as the protease and allowed a maximum of two missed cleavages and semi-specific digestion. Carbamidomethylation of cysteine (+57.02 Da) was set as fixed modification and methionine oxidation (+15.99 Da), lysine lactylation (+72.02 Da) and acetylation at protein N termini (+42.02 Da) were set as variable modifications. Peptide-spectrum matches (PSMs) were filtered to 1% FDR employing a target-decoy database search approach. The modified peptides reached an AScore>20^65^ was assigned as lactylated candidates. Further, MS/MS spectra of such peptides were classified as true hits on condition that the signature CycIm ion must be present in the examined spectra.

### Wet-lab strategy for lactylation mining

#### Sample preparation

To identify novel lactylation sites, cells were treated with 25 mM sodium lactate for 24 h, washed with cold PBS for three times, harvested and lysed in RIPA lysis buffer (Beyotime, cat. no. P1003B) with protease and phosphatase inhibitor cocktail (ApexBio, cat. no. K1007 and K1013). Then, methanol, chloroform and water were added to the lysate at a ratio of 4:1:3:1 by volume. Precipitated protein was collected by centrifugation at 12,000 rpm for 10 min and washed twice with methanol, followed by re-solubilized by 8 M urea in 25 mM ammonium bicarbonate solution. Proteins were then reduced by 10 mM dithiothreitol (DTT) at 56 ℃ for 30 min and alkylated by 40 mM iodoacetamide (IAM) at 25 ℃ for 20 min in dark. Additional DTT was added to react with excess IAM at 25 ℃ for 10 min. Subsequently, the mixtures were added with 25 mM ammonium bicarbonate to dilute urea to 1 M, followed by digestion with sequencing-grade trypsin (Promega, cat. no. V5111) at an enzyme/protein ratio of 1:50 (w/w) overnight at 37 ℃. The lactylated peptides were enriched by immunoprecipitation according to literature^3^. Briefly, protein A agarose beads (Invitrogen, cat. no. 15918014) were washed with ETN buffer (1 mM EDTA, 50 mM Tris-HCl (pH 8.0) and 100 mM NaCl) for three times, blocked with 5% BSA and conjugated with anti-lactylation antibody at 4 ℃ overnight. Then, the digested peptides were resolved with ETN buffer and incubated with the prepared antibody-conjugated beads with rotary shaking at 4 ℃ for 6 h. The beads were washed for three times with ETN buffer and the beads-bounded peptides were eluted with 1% trifluoroacetic acid in 40% ACN. The eluted peptides were evaporated to dryness and stored at −80 ℃ prior to analysis.

#### Lactylproteome analysis

The digested cell lysates were analyzed by an Orbitrap Eclipse Tribrid mass spectrometer equipped with an EASY-nano LC 1200LC system (Thermo Fisher Scientific). The mobile phase consisted of solvent A (0.1% FA in water) and solvent B (ACN/Water, 8:2, v/v). The flow rate was set at 300 nL/min. Peptides were analyzed using an Acclaim PepMap RSLC column (75 μm × 250 mm, Thermo Fisher Scientific) with a 90-min chromatography gradient: 0-5 min, 3-8% phase B; 5-6 0min, 8-28% phase B; 60-75 min, 28-38% phase B; 75-80 min, 38-100% B; 80-90 min, 100% B. For MS data acquisition, MS1 spectra were collected at the *m/z* range of 350-1,800 at a resolution of 120,000 on the Orbitrap with a maximum AGC value of 4e^5^. For MS2 acquisition, fragmentation was conducted by HCD with the CE value set at 32% after optimization. MS2 spectra were collected with the first mass set at *m/z* 110 at a resolution of 30,000 on the Orbitrap with a maximum AGC of 5e^4^.

#### Data analysis

Lactylproteome data were searched using similar parameters as dry-lab analysis with PEAKS Studio XPro against the UniProt human proteome database (UniProt_Human_reviewed_15-5-2023.fasta) and UniProt_Mouse_reviewed_11-6-2023.fasta. Specifically, mass tolerance was set to 10 ppm for precursors and 0.02 Da for fragment ions, and FDR was filtered to 1% for PSMs. No additional CycIm ion filtering is required.

### Phylogenetic analysis

Protein sequences of ALDOA from each species were aligned with the Molecular Evolutionary Genetics Analysis X (MEGA X) software^66^. The phylogenetic tree was constructed by MEGA X using the Neighbor-Joining method with 1,000 bootstrap replicates. The tree is drawn to scale, with branch lengths in the same units as those of the evolutionary distances used to infer the phylogenetic tree. The evolutionary distances were computed using the Poisson correction method. Scale bar indicates the number of amino acid substitutions per site.

### KlacRS selection and engineering

The focused mutant library of pBK-*Mm*PylRS was constructed and transformed into DH10B cells containing pREP positive selection plasmid via electroporation. The cells were recovered with 1 mL pre-warmed SOC medium and shaken vigorously at 37 ℃ for 1 h. The recovered cells were plated on LB agar plate containing 50 μg/mL kanamycin (Kan), 25 μg/mL tetracycline (Tet), 100 μg/mL chloramphenicol (Cm), 0.02% L-arabinose (Ara) and 1 mM Klac. The plate was cultivated at 37℃ for 48 h. Colonies with strong green fluorescence were picked and replicated on LB agar plates containing 50 μg/mL Kan, 25 μg/mL Tet, 100 μg/mL Cm and 0.2% Ara with or without 1 mM Klac. A clone exhibiting Klac-dependent growth and fluorescence was named as pBK-*Mm*KlacRS2 and Sanger sequenced. To compare its incorporation efficiency with other KlacRS variants, *Mm*KlacRS2 was then cloned into pEvol vector by homologous recombination following the manufacturer’s instructions (Vazyme, cat. no. C112).

To construct the chimeric KlacRS clones, namely pEvol-chKlacRS-WT and pEvol-chKlacRS-IPYE, we fused residues 1-149 of either wild type *Mb*PylRS (*Mb*PylRS-WT) or its activated mutant (*Mb*PylRS-IPYE) with residues 185-454 of *Mm*KlacRS1 by overlapping PCR and then cloned into pEvol vectors by homologous recombination. The DNA sequences of KlacRS variants are shown in **Supplementary Table 5** and the primers used are listed in **Supplementary Table 6**.

### Incorporation efficiency of KlacRS variants

The plasmids pEvol-*Mm*KlacRS1, pEvol-*Mm*KlacRS2, pEvol-chKlacRS-WT, and pEvol-chKlacRS-IPYE were co-transformed into DH10B cells with pBad-EGFP-39TAG reporter, respectively. The transformants were grown in 2 mL LB containing 100 μg/mL ampicillin (Amp) and 50 μg/mL Cm at 37 ℃ until OD600 reached 0.6, followed by adding 0.2% Ara with or without 1 mM Klac. After grown at 30 ℃ for another 16 h, cells were collected by centrifugation, washed once with PBS, and then resuspended in PBS. The relative fluorescence units (RFU) of cells were determined by dividing the absolute fluorescence by the OD600 readings of each sample using BioTek Synergy H1 microplate reader (Agilent, Santa Clara, CA, USA) to compare the incorporation efficiency of above KlacRS variants. The wild type EGFP cultured under the same conditions was used as a control.

### Cell transfection, Klac incorporation and fluorescence imaging

For siRNA transfection, siCtrl and siALDOA were purchased from RiboBio (Guangzhou, China). Approximately 3×10^5^ HEK293T cells were seeded into 6-well plates and transfected with 20 nM siRNA using 5 μL lipofectamine^TM^ RNAiMAX reagent (Thermo Fisher Scientific, cat. no. 13778150) for 48 h according to the manufacturer’s instructions. The efficiency of silencing was confirmed by immunoblotting.

To genetically encode Klac into target proteins, approximately 3×10^5^ HEK293T cells were seeded into 6-well plates and cultured for 24 h. Plasmids pCMV-EGFP-Y39TAG and pcDNA-mCherry-TAG-EGFP were co-transfected with the pNEU-*Mm*KlacRS1 plasmid, respectively, using PolyJet transfection reagents according to the manufacturer’s instructions. Then, the cultured media were replaced with fresh complete medium with or without indicated concentration of Klac at 18 h post-transfection and cells were cultured for another 24 or 48 h.

The transfected cells were examined by fluorescence imaging using Leica DMI 3000B light microscope (Leica, Wetzlar, Germany) and relative fluorescence intensities between cells cultured with or without Klac were calculated by ImageJ (National Institutes of Health, V1.51). Following the same procedures, the plasmid pCMV-ALDOA-K147TAG were co-transfected with pNEU-*Mm*KlacRS1 and subjected to immunoblotting analysis.

### Flow cytometry analysis

Approximately 3×10^5^ HEK293T cells were seeded into 6-well plates and cultured for 24 h. Plasmids pcDNA-sfGFP-Y150TAG or pCMV-EGFP-Y39TAG were co-transfected with pNEU-*Mm*KlacRS1 and cultured in the presence or absence of the indicated concentration of Klac for additional 24 h and 48 h. Cells were washed with cold PBS, harvested by centrifugation and resuspended in PBS. The fluorescence of cells was measured by CytoFlex flow cytometer using 405/488 nm lasers (Beckmann Coulter, CA, USA).

### Immunoblotting

Cells were lysed in RIPA lysis buffer supplemented with protease and phosphatase inhibitor cocktail. Nuclear and Cytoplasmic Protein Extraction Kits (Beyotime, cat. no. P0027) were used for separating cytoplasmic and nuclear proteins. Protein concentrations were first determined by the bicinchoninic acid (BCA) assay (Beyotime, cat. no. P0011). Then, the lysates were diluted by 4×XT Sample Buffer (Bio-rad, cat. no.1610791), heated to 100 ℃ for 5 min, cooled and separated by 10% SDS-PAGE gels. Proteins were then transferred onto polyvinylidene difluride (PVDF) membranes (Bio-rad, cat. no.1620177), blocked with 5% non-fat dry milk in Tris-buffered saline with 0.1% Tween 20 detergent (TBST) and incubated with primary antibodies at 4 ℃ overnight. The membranes were subsequently washed five times with TBST and incubated with horseradish peroxidase (HRP)-conjugated secondary antibody for 1 h at 37 ℃. The immunoblotted bands were detected by the addition of HRP substrate (Bio-rad, cat.no. 1705601), captured on a ChemiDoc XRS+ system (Bio-rad, Hercules, USA) and analyzed by ImageLab software. The primary antibodies used in this study include the antibody against ALDOA (Proteintech, 11217-1-AP, 1:1000), His_6_-tag (Cell Signaling Technology, 2365S, 1:1000), Flag-tag (Cell Signaling Technology, 14793S, 1:1000), lysine lactylation (PTM Bio Inc, PTM-1401RM, 1:2000), H4-K9-lactylation (PTM Bio Inc, PTM-1415RM, 1:1000), H4-K13-lactylation (PTM Bio Inc, PTM-1411RM, 1:1000), β-tubulin (Proteintech, 10068-1-AP, 1:1000) and β-actin (Proteintech, 66009-1-Ig, 1:20000).

### ALDOA activity assay

Enzymatic activity of ALDOA-WT and ALDOA-147Klac were assessed using an aldolase activity colorimetric assay kit (Biovision, cat. no. K665-100) according to the manufacturer’s instructions as previously reported^3^. Briefly, cells were collected and homogenized with ice-cold aldolase assay buffer and kept on ice for 10 min. Cell lysates were centrifuged and the supernatant was collected. The supernatants were then diluted with the assay buffer to prepare the sample solution. Different volumes of NADH standard were used to generate a NADH standard curve. The reaction solution was prepared by mixing the aldolase enzyme mix, aldolase developer, aldolase substrate and aldolase assay buffer. The reaction was initiated by the addition of 50 μL reaction solution to 50 μL sample solution and standard solution, respectively. ALDOA activity was measured photometrically by monitoring the absorbance at 450 nm in kinetic mode for 30 min with a 1-min interval at 37 ℃. Data were normalized to the protein concentrations as determined by BCA assay for each sample.

### Metabolomic analysis of glycolytic metabolites

Cells were washed twice with cold PBS and intracellular metabolites were extracted by the addition of 1 mL extraction solvent that consisted of methanol/ACN/water (40:40:20, v/v/v, pre-cooled at −20 ℃) and 0.5 μg/mL 4-chloro-phenylalanine as an internal standard (IS). Following co-incubation for 30 min at −20 ℃, cells were scraped off and the mixtures were transferred to 1.5 mL centrifugation tubes, followed by centrifugation at 18,000 g for 10 min at 4 ℃. The resultant supernatant was collected and evaporated to dryness, then analyzed by LC/MS.

Metabolites were analyzed using an Agilent UHPLC 1290 (Agilent Technologies, Santa Clara, CA USA) coupled to an Agilent 6546 Q/TOF mass spectrometer. Chromatographic separation was achieved using an XBridge Amide HPLC column (100 × 4.6 mm, 3.5 μm, Waters). The mobile phase consisted of solvent A (10 mM ammonium acetate in water, pH adjusted to 9.0 by ammonia) and solvent B (ACN). The gradient was set as follows: 0-1 min, 85% B; 1-16 min, 85-30% B; 16-18 min, 30% B; 18-18.5 min, 30-85% B; 18.5-30 min, 85% B. The flow rate was set at 0.4 mL/min and the column temperature was set as 40 ℃. Following separation, the Agilent 6546 Q-TOF was operated in the negative mode (ESI-) for metabolite detection. The following parameters were used: liquid nebulizer, 45 psi; nitrogen drying gas, 8 L/min; drying gas temperature, 320; ESI capillary voltage, −3500 V; MS scan, *m/z* 50-1100; fragmentor voltage, 80 V. The Agilent Masshunter Workstation Data Acquisition was used for data acquisition and Agilent Masshunter Qualitative Analysis for data analysis. Metabolite identification was performed by matching the retention time, the precursor *m/z* and the fragment ions against analytical standards.

### Metabolic measurement by Seahorse

For Seahorse analysis, ECAR of cells was measured using the Seahorse XF Glycolysis Stress Test Kit (Agilent Technologies, Palo Alto, CA, USA) on an XFe96 extracellular flux analyzer (Seahorse Bioscience, Billerica, MA, USA). Briefly, 1×10^5^ HEK293T cells were seeded onto Seahorse XF96 microplates and cultured overnight. Then, the cultured media were replaced with the assay medium (XF base medium containing 2 mM glutamine, pH=7.4), and cells were equilibrated at 37 ℃ in a CO_2_-free incubator for 1 h. After baseline measurement, the standard Glycolysis Stress test was carried out with the stepwise injection of 10 mM glucose, 3 uM oligomycin and 50 mM 2-deoxy-D-glucose (2-DG). Metabolic parameters were collected and analyzed using the XF Wave software (Agilent, Palo Alto, CA, USA). Data were normalized to the protein concentrations as determined by BCA assay for each well.

### DARTS assay

HEK293T cells expressing Flag-tagged ALDOA-WT and ALDOA-147Klac were lysed in M-PER buffer (Thermo Scientific, cat. no. 78501) supplemented with protease and phosphatase inhibitor cocktail. Protein concentrations were determined by BCA assay. Cell lysates were diluted with TNC buffer (50 mM Tris-HCl at pH 8.0, 50 mM NaCl and 10 mM CaCl_2_) to 2 mg/mL and then divided into 7 aliquots of 40 μL, respectively. Different concentrations of Pronase (Roche, cat. no. 10165921001) were added to aliquoted lysates followed by incubation at 25 ℃ for 30 min. Proteolysis was quenched by the addition of 4×XT Sample Buffer and boiling for 10 min. Samples were then subjected to immunoblotting analysis using antibodies against the Flag tag and β-actin. The resultant band intensities of the Flag-tagged ALDOA were normalized to the intensity of the Pronase-untreated group, and fitted to the Boltzmann sigmoid equation using Prism v.8.0.1 (GraphPad software).

### CETSA

HEK293T cells expressing ALDOA-WT and ALDOA-147Klac were aliquoted, followed by heating at designed temperatures ranging from 56 to 77.5 ℃ for 3 min in the Thermal Cycler (2720 Thermal Cycler, Applied Biosystems, CA, USA) and cooling down at room temperature for another 3 min. All samples were then subjected to three freeze-thaw cycles and centrifuged at 20,000 g for 20 min. The soluble fractions were diluted by 4×XT Sample Buffer, heated to 100 ℃ for 5 min, and subjected to immunoblotting analysis. The resultant band intensities of the Flag-tagged ALDOA were normalized to the intensity of the unheated group, and fitted to the Boltzmann sigmoid equation using Prism v.8.0.1 (GraphPad software)^67^. The melting temperature (T_m_) is defined as the temperature at which a 50% reduction in signal (soluble protein) is observed.

### Confocal imaging

HEK293T cells expressing ALDOA-WT and ALDOA-147Klac were plated on 35 mm glass-bottom dishes (Nest, Jiangsu, China). Cells were washed with PBS, fixed in 4% paraformaldehyde for 15 min at room temperature in dark, and permeabilized with 0.1% Triton X-100 for 10 min. Cells were then incubated with Hoechst 33258 (KeyGEN, cat. no. KGA1802) for 5 min at room temperature in dark and washed with PBS. Images of cells were acquired using a Zeiss 700 confocal microscope (Zeiss, Jena, Germany).

### RNA-seq analysis

Total RNA was extracted using TRIzol reagent (Thermo Scientific, catalog no. 15596026), and RNA-seq was performed by Annoroad Gene Technology (Beijing, China). Briefly, the integrity and concentration of RNA were analyzed using an Agilent RNA 6000 Nano Kit (Agilent, cat. no. 5067-1511) and Agilent 2100 Bioanalyzer (Agilent Technologies). RNA-seq was performed using an Illumina Nova 6000 platform (Illumina), and the Novaseq Control Software (v.1.7.5) was used for data collection. Raw sequenced reads were filtered to achieve high-quality reads and then mapped to the human genome (GRCh38.100.chr) using HISAT2. Differentially expressed genes (DEGs) were analyzed using the R package DESeq2 with a cutoff of FC > 1.5 or <0.67 and P value <0.05 (n=4 biologically independent samples/group). DEGs were subjected to KEGG pathway analysis and GO functional annotation including molecular function, biological process and cellular component by Metascape^68^ (https://metascape.org/). A setting of group P value <0.01 and the inclusion of at least three genes in each group was used for filtering.

### RNA extraction and RT-qPCR analysis

Cells were harvested and washed with PBS twice. Total RNA was isolated using FreeZol Reagent (Vazyme, cat. no. R711-01) according to the manufacturer’s instruction. Approximately 200 ng of total RNA was reverse transcribed into cDNA using Hiscript II RT SuperMix (Vazyme, cat. no. R222-01). RT-qPCR analysis of target genes was then performed using SYBR qPCR Master Mix (Vazyme, cat. no. Q341-02) on a real-time PCR cycler (StepOne Plus, Applied Biosystems, CA, USA). The endogenous β-tubulin gene was used as a reference to normalize the expression levels of target genes. The primer sequences used in this study are listed in **Supplementary Table 6**.

### Co-IP analysis of ALDOA interactome

Cells were harvested and lysed in 0.5% NP-40 lysis buffer (Beyotime, cat. no. P0013F) with protease inhibitor cocktail, phosphatase inhibitor cocktail and super nuclease (Beyotime, cat. no. D7271) followed by centrifugation at 18,000 g for 10 min. Protein concentration was determined by BCA assay, and approximately 1 mg of protein lysates were incubated with anti-Flag magnetic beads (Selleck, cat. no. B26101) for 4 h at room temperature for immunoprecipitation. For immunoblotting, the beads-bound proteins were eluted with SDS loading buffer and then heated at 100 ℃ for 5 min. For the ALDOA interactome analysis, the beads-bound proteins were then washed with PBS for three times and incubated with 8 M urea in 25 mM ammonium bicarbonate solution for denaturation. Proteins were reduced by 10 mM DTT at 56 ℃ for 30 min and alkylated by 40 mM IAM at 25 ℃ for 20 min in dark. Additional DTT was added to react with excess IAM at 25 ℃ for 10 min. Subsequently, the mixtures were added with 25 mM ammonium bicarbonate for diluting urea to 1 M, followed by digestion with trypsin at an enzyme/protein ratio of 1:50 (w/w) overnight at 37 ℃. Samples were then acidified, desalted with C_18_ Zip-tips, evaporated to dryness, followed by analyzed using the Orbitrap Eclipse Tribrid mass spectrometer with the EASY-nano LC with a 90-min chromatography gradient as previously described. Data collection and analysis were also performed as previously described.

### Functional annotation of ALDOA interactome

The identified interactome of ALDOA-WT and ALDOA-147Klac was annotated to GO terms and KEGG pathway analysis. Specifically, GO CC annotation was performed using the DAVID bioinformatics website (https://david.ncifcrf.gov/). For GO MF annotation and KEGG pathway analysis, Metascape (http://metascape.org/) was used to perform the functional enrichment analysis. For GO BP terms, Metascape was employed to perform the enrichment analysis and Cytoscape was employed to visualize the annotation network. A group P value <0.05 for CC annotation and a group P value <0.01 for GO BP, MF annotation and KEGG pathway analysis. At least three genes were included in each group for filtering.

## Data availability

The Meltome Atlas^22^, the deep proteome atlas of 29 healthy human tissues^23^, the proteome of kingdoms of life^24^ and an affinity-enriched lactylatome of Western flower thrips^25^ were accessed through the PeoteomeXchange Consortium (https://proteomecentral.proteomexchange.org) with the dataset identifier PXD011929, PXD010154, PXD014877 and PXD030799, respectively. Experimental data collected in this study can be accessed through the Consortium via the iProX partner repository^69^ with the dataset identifier PXD028488 and PXD048127. RNA sequencing data presented in this study are in the NCBI GEO repository under the accession number GSE252184.

