## Supplementary Figures for "Genetic code expansion reveals site-specific lactylation in living cells reshapes protein function"

**a** **Summary of ALDOA-K147 Occupancy**

| Cell line | K562-R1 | K562-R2 | Jurkat-R2 | A549-R2 | Lung fibro.-R1 | HL60-R2 | Colon sph.-R1 | Colon sph.-R2 | HaCat-R1 | HAOEC-R1 | HAOEC-R2 | HEK293T-R2 |
|---|---|---|---|---|---|---|---|---|---|---|---|---|
| Occupancy (%) | 4.78 | 3.03 | 9.11 | 50.33 | 33.26 | 13.93 | 6.84 | 13.18 | 0.1 | 0.8 | 0.22 | 0.17 |

**b**

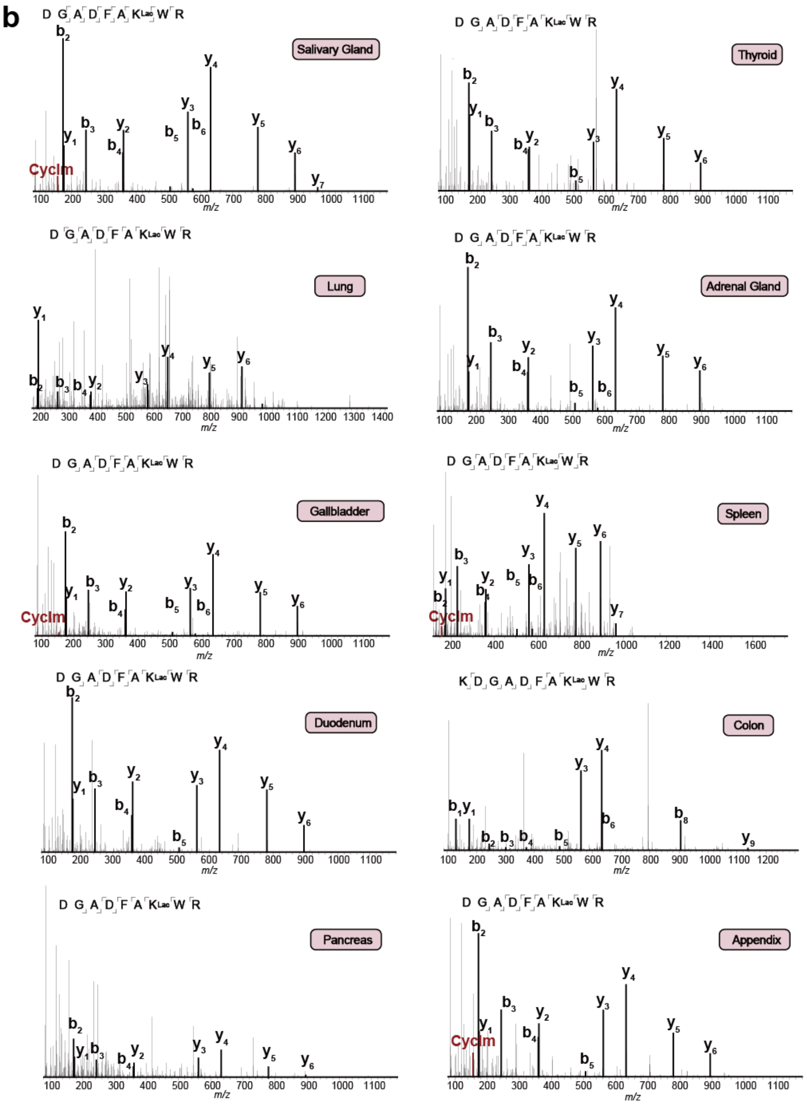

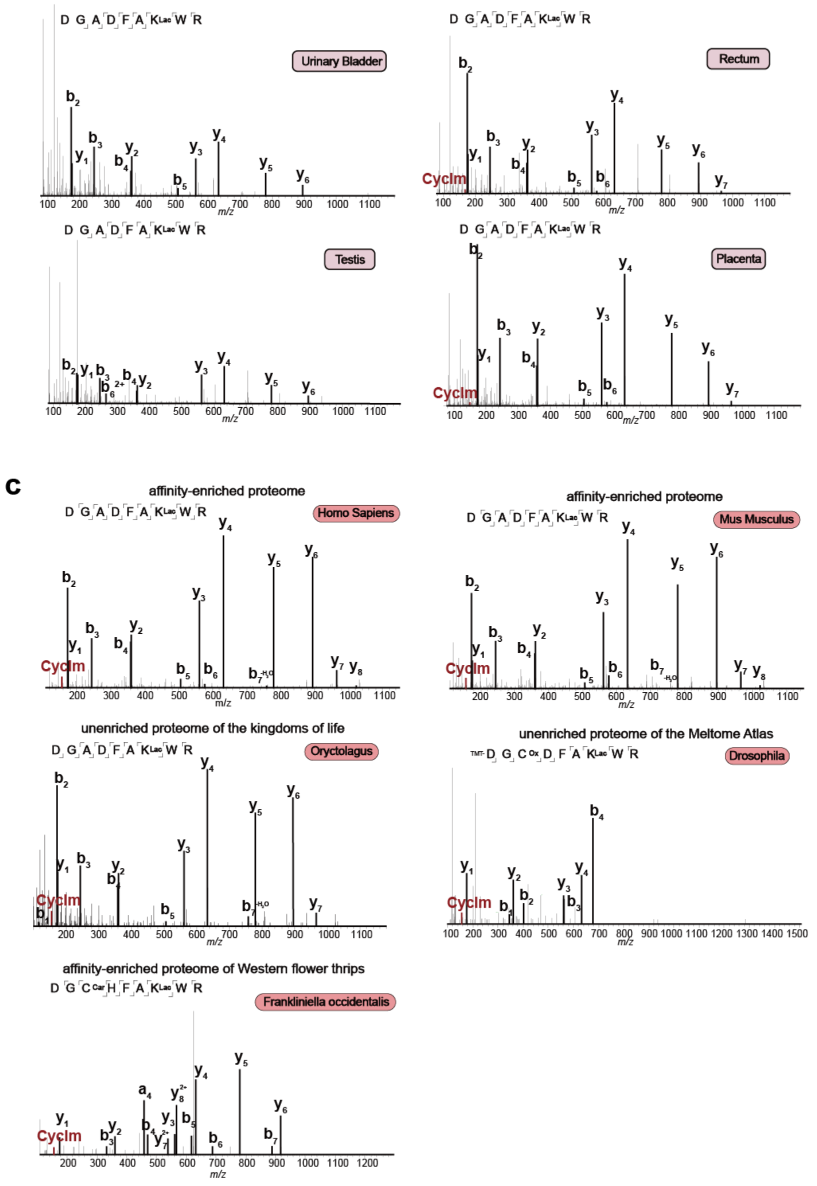


**Supplementary Figure 1. Proteomics mining pinpoints functionally important lactylation on ALDOA.**

(a) Re-presentation of previously estimated lactylation occupancy for K147 of ALDOA[1] using human cell proteome data retrieved from the Meltome Altas (PXD011929).

(b) Representative MS/MS spectra leading to the assignment of lactylation on K147 of ALDOA in various human tissues retrieved from the deep proteome atlas of 29 human healthy tissues (PXD010154), including salivary gland, thyroid, lung, adrenal gland, gallbladder, spleen, duodenum, colon, pancreas, appendix, urinary bladder, rectum, testis and placenta.

(c) Representative MS/MS spectra leading to the assignment of lactylation on K147 of ALDOA in the 5 species, including *Homo sapiens* (HEK293T) and *Mus musculus* (RAW264.7) from our affinity-enriched lactylproteome, *Oryctolagus cuniculus* from the re-analyzed unenriched proteome of the kingdoms of life (PXD014877), *Drosophila* from the re-analyzed proteome retrieved from the Meltome Altas (PXD011929) and *Frankliniella occidentalis* from an affinity-enriched lactylproteome of Western flower thrips (PXD030799).

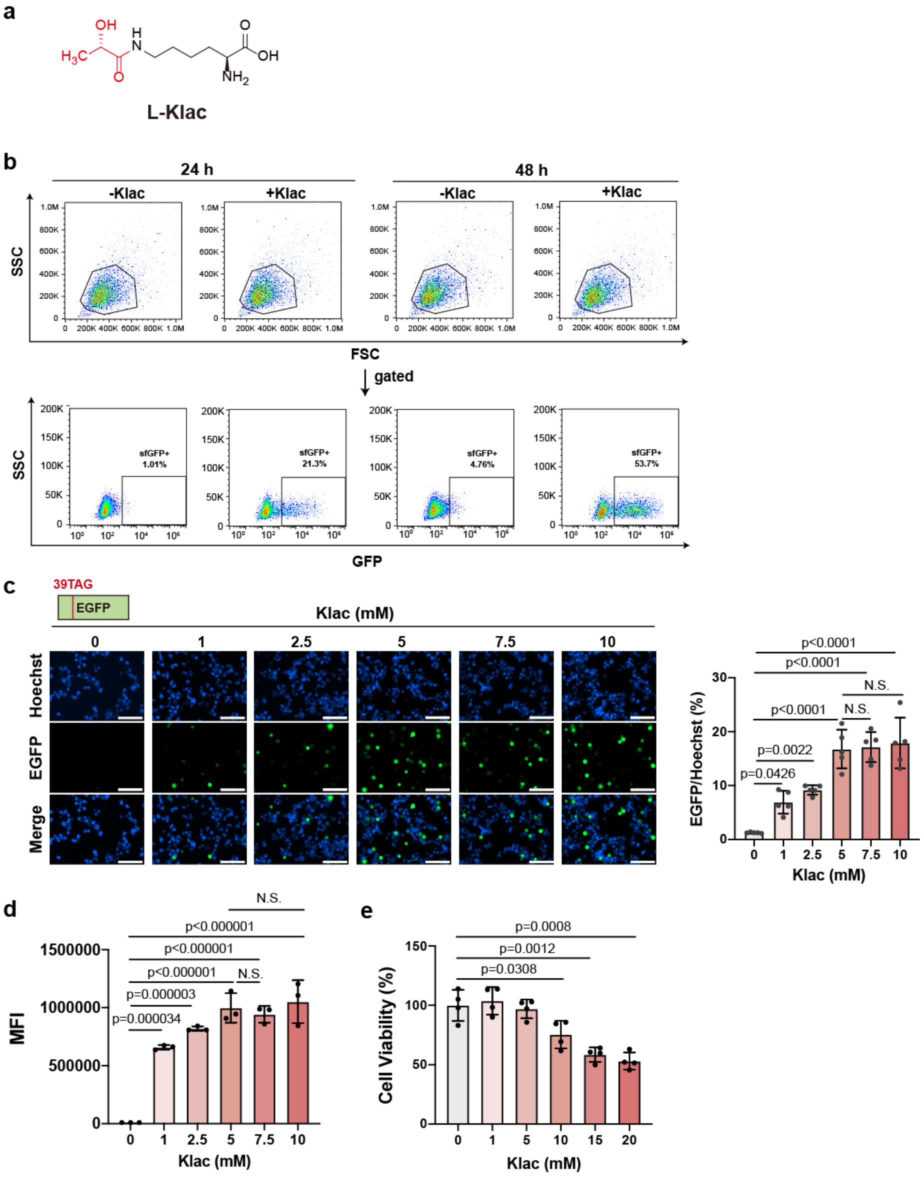


**Supplementary Figure 2. Site-specific incorporation of lactylation in living cells with genetic code expansion**

(a) Structure of synthesized L-Klac.

(b) Gating strategy to sort HEK293T cells co-transfected with KlacRS1/$tRNA^{Pyl}_{CUA}$ pair and sfGFP-150TAG plasmids treated with or without Klac (1 mM) for 24 h and 48 h as specified in **Fig. 2 c**.

(c) Representative images of HEK293T cells co-transfected with KlacRS1/$tRNA^{Pyl}_{CUA}$ pair and EGFP-39TAG plasmids treated with indicated concentrations of Klac for 48 h. The nucleus was marked with Hoechst (blue). Data represent the mean ± S.D. (n=5 biological replicates/group) and the p value was calculated by one-way ANOVA.

(d) Flow cytometry analysis of HEK293T cells co-transfected with KlacRS1/$tRNA^{Pyl}_{CUA}$ pair and EGFP-39TAG plasmids treated with indicated concentrations of Klac for 48 h. Data represent the mean ± S.D. (n=3 biological replicates/group) and the p value was calculated by one-way ANOVA.

(e) Cell viability of HEK293T cells treated with indicated concentrations of Klac for 48 h. Data represent the mean ± S.D. (n=4 biological replicates/group) and the p value was calculated by one-way ANOVA.

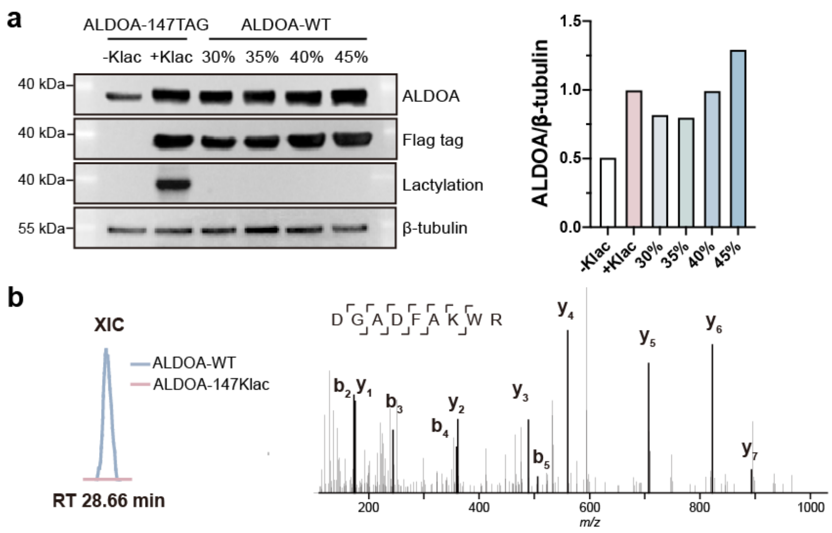


**Supplementary Figure 3. Introducing site-specific lactylation on K147 of ALDOA in living cells.**

(a) Immunoblotting analysis of HEK293T cells co-transfected with KlacRS1/tRNA$^{Pyl}_{CUA}$ pair and ALDOA-147TAG plasmids with or without Klac (5 mM, 48 h), and those transfected with different amounts of ALDOA-WT plasmids. Cells expressing ALDOA-147Klac and ALDOA-WT at similar levels (Lane 2 and 5) were used for following functional studies. Left, representative images of western blots. Right, quantification of ALDOA levels normalized to ß-tubulin.

(b) Identification of non-lactylated K147 peptides in cells expressing Flag-tagged ALDOA-WT but not in those expressing ALDOA-147Klac. HEK293T cells transfected with ALDOA-WT or co-transfected with KlacRS1/tRNA$^{Pyl}_{CUA}$ pair and ALDOA-147TAG treated with Klac (5 mM, 48 h), followed by immunoprecipitation with anti-Flag antibody and bottom-up proteomic analysis. Left, XIC of non-lactylated K147 peptide. Right, representative MS/MS spectrum of non-lactylated K147 peptide.

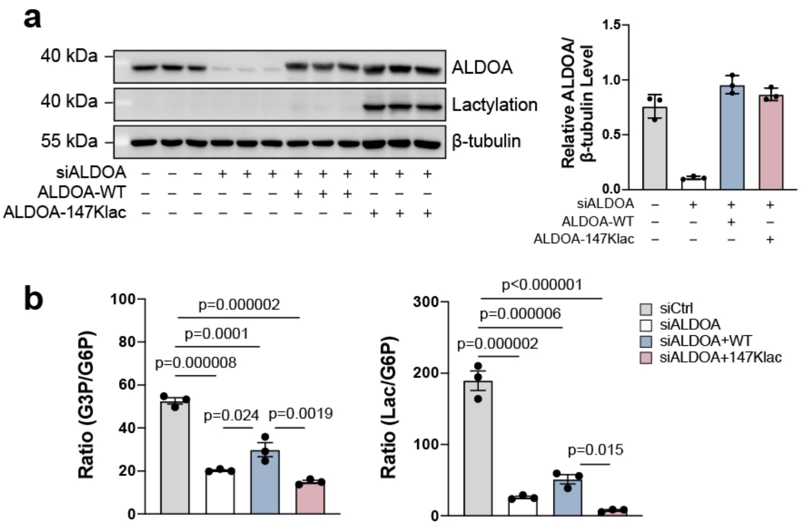


**Supplementary Figure 4. Lactylation on ALDOA-K147 abolished enzyme activity and impaired glycolytic flux.**

(a) HEK293T cells expressing ALDOA-WT or ALDOA-147Klac after knocking down the endogenous ALDOA were used to determine the changes in enzymatic activity of ALDOA after lactylation. Left, representative images of western blots. Right, quantification of ALDOA levels normalized to ß-tubulin. Data represent the mean ± S.D. (n=3 biological replicates/group) and the p value was calculated by one-way ANOVA.

(b) Abundance ratios of G3P/G6P and Lac/G6P in the siCtrl, siALDOA, siALDOA+WT and siALDOA+147Klac groups. Data represent the mean ± S.D. (n=3 biological replicates/group) and the p value was calculated by one-way ANOVA.

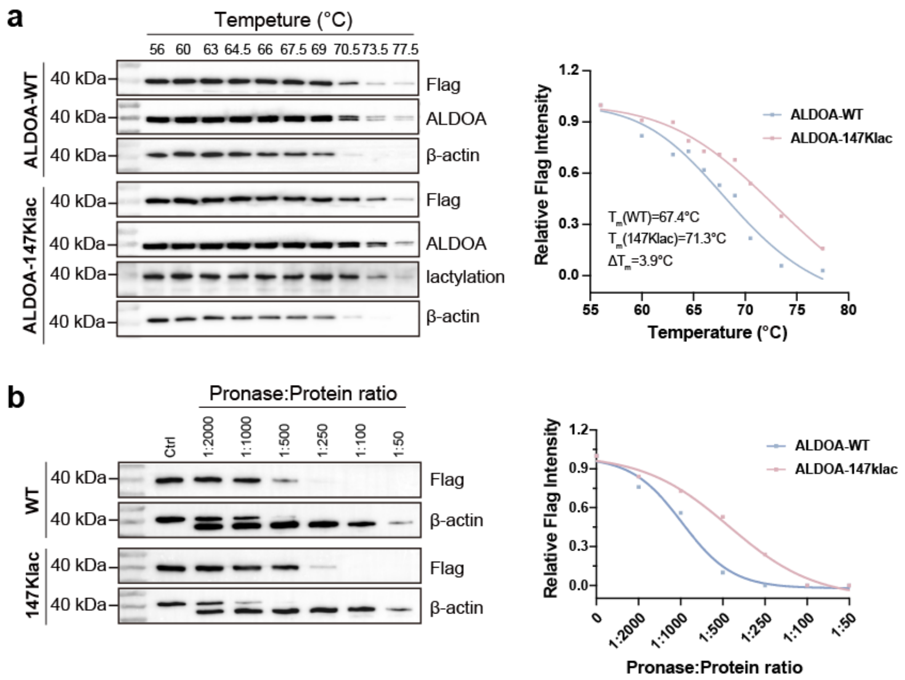


**Supplementary Figure 5. Site-specific lactylation modulated ALDOA stability**

(a) Immunoblotting-based CETSA using HEK293T cells expressing ALDOA-WT, via transfection with ALDOA-WT plasmid, and cells expressing ALDOA-147Klac, via co-transfection with KlacRS1/ $tRNA^{Pyl}_{CUA}$ pair and ALDOA-147TAG plasmids and cultured with Klac (5 mM, 48 h). Left, representative images of western blots. Right, band intensity profiles. Melting temperature ($T_m$) is defined as the temperature at which a reduction of 50% signal (soluble protein) is observed. Representative immunoblots and melting curves were shown and experiments were repeated three times with consistent results.

(b) Immunoblotting-based DARTS assay using HEK293T cells in (a). Left, representative images of western blots. Right, band intensity profiles. Experiments were repeated (n = 3 biologically independent samples).

**a**

| Location | Probability of Location |
|---|---|
| Cytoplasm | 90.6% |
| Nucleus | 9.2% |
| Mitochondrion | 0.2% |

**b**

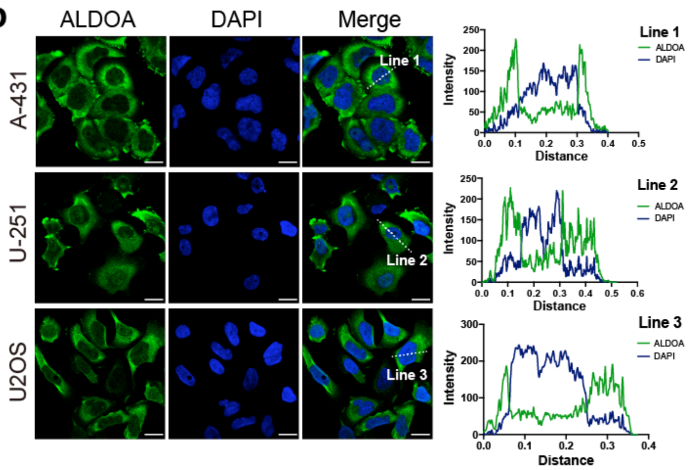


**Supplementary Figure 6. Subcellular localization analyses of ALDOA-WT show primary localization in cytoplasm.**

(a) Subcellular localization of ALDOA was predicted via YLoc[2] analysis with a confidence of 0.82 (https://abi-services.cs.uni-tuebingen.de/yloc/webloc.cgi).

(b) Subcellular localization of ALDOA in human cell lines A-431, U-251 and U2OS by immunofluorescence staining analysis retrieved from the Human Protein Altas[3] (https://www.proteinatlas.org/ENSG00000149925-ALDOA/subcellular). ALDOA was labeled with the fluorescent protein EGFP (green) and the nucleus was labeled with DAPI (blue). Left, representative images showing that ALDOA was primarily localized in the cytoplasm. Right, fluorescence intensity profiles across the lines indicated on the left. Scale bar, 20 μm.

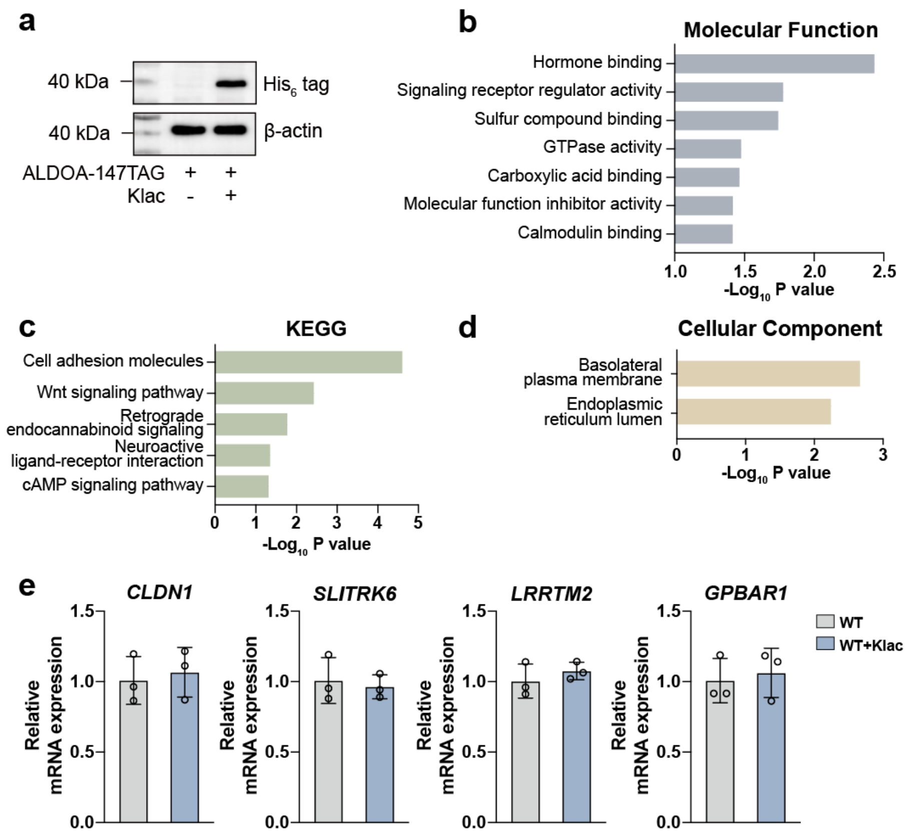


**Supplementary Figure 7. Lactylation on ALDOA induced transcriptional changes in living cells.**

(a) HEK293T cells co-transfected with KlacRS1/tRNA$^{Pyl}_{CUA}$ pair and ALDOA-147TAG plasmids treated with or without Klac (5 mM, 48 h) were used for RNA-seq analysis.

(b-d) Bar chart of the GO MF analysis (b), KEGG pathway analysis (c) and the CC analysis (d) for the differentially regulated genes using Metascape with the Benjamini-Hochberg correction algorithm.

(e) RT-qPCR analysis of *CLDN1*, *SLITRK6*, *LRRTM2* and *GPBAR1* expression in cells transfected with ALDOA-WT with and without Klac (5 mM, 48 h). The endogenous β-tubulin gene was used as the internal control for normalizing target gene expression changes. Data represent the mean ± S.D. (n=3 biological replicates/group) and the p value was calculated by one-way ANOVA.

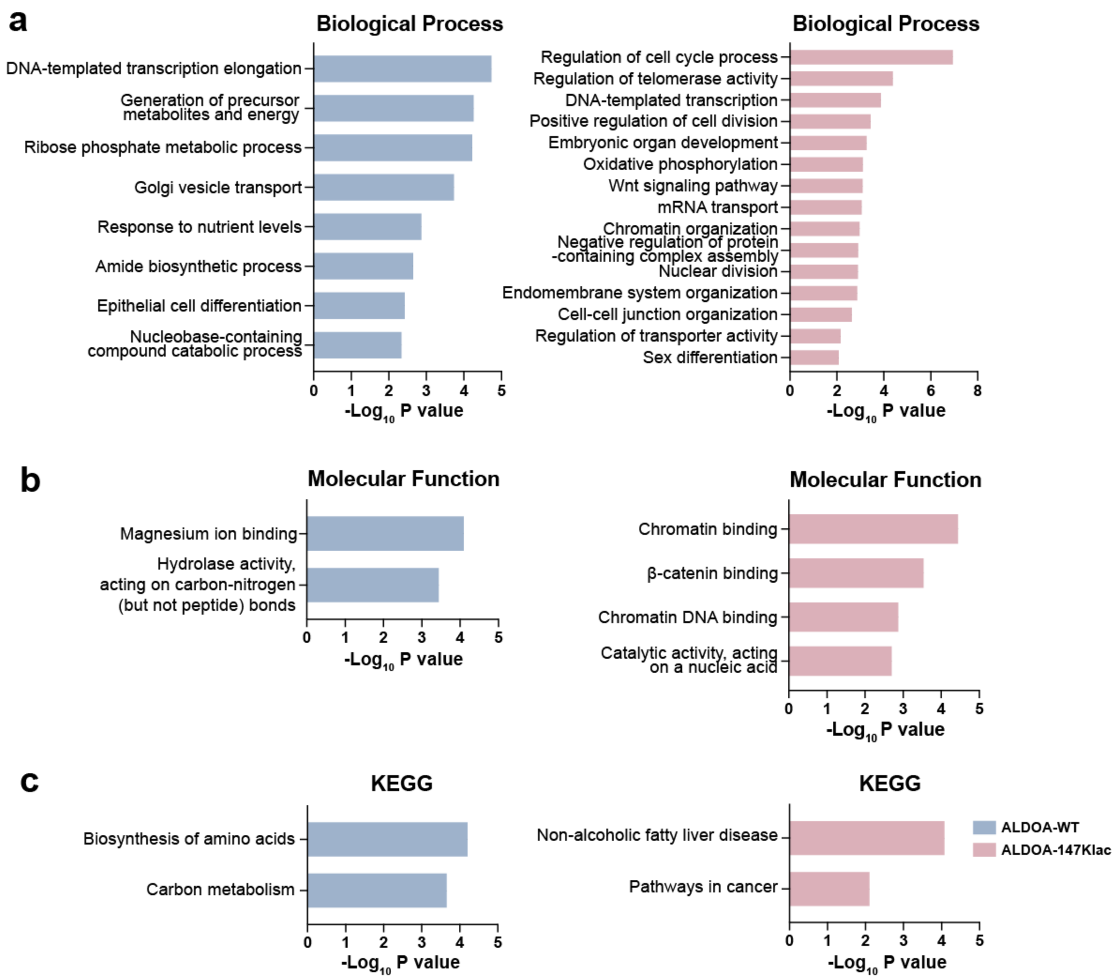


**Supplementary Figure 8. ALDOA recruited different interacting partners after lactylation**

(a-c) Bar chart of the GO BP analysis (a), MF analysis (b), and the KEGG pathway analysis (c) for the interacting proteins of ALDOA-WT and ALDOA-147Klac using Metascope with the Benjamini-Hochberg correction algorithm.

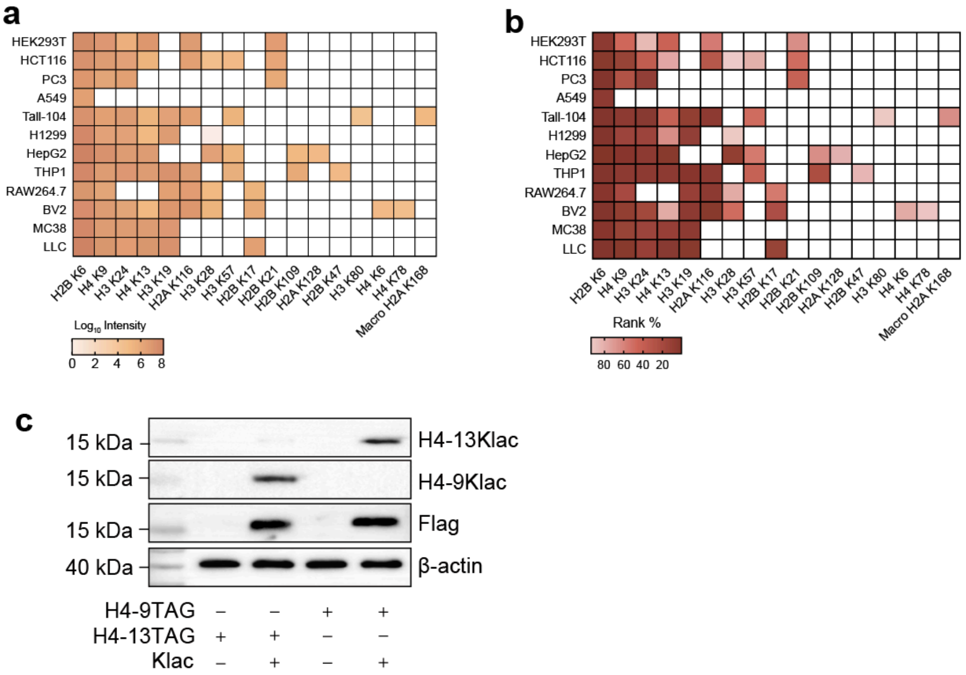


**Supplementary Figure 9. Site-specifically encoding histone lactylation in living cells**

(a-b) The ion intensity (a) and the ranking (b) of the lactylated sites identified on histone in our affinity-enriched lactylproteome of representative human and mouse cell lines. Cells were treated with 25 mM lactate for 24 h to stimulate lactylation.

(c) Validation of site-specific incorporation of lactylation at K9 and K13 of histone H4, respectively. HEK293T cells co-transfected with KlacRS1/$tRNA^{Pyl}_{CUA}$ pair and H4-9TAG or H4-13TAG treated with or without Klac (5 mM, 48 h). Full-length of histone H4 was estimated using anti-Flag antibody. Genetically engineered histone H4 lactylation was detected using anti-H4-9Klac and anti-H4-13Klac antibodies, respectively.

### Supplementary Tables

**Table 1.** Summary of the lactylated peptides identified from the 12 cell lines.

**Table 2.** Summary of the peptides carrying ALDOA-147Klac identified from 15 healthy tissues retrieved from the deep proteome atlas of 29 healthy human tissues.

**Table 3.** Lactylation occupancy for K147 of ALDOA in 15 healthy tissues retrieved from the deep proteome atlas of 29 healthy human tissues.

**Table 4.** Summary of the peptides carrying ALDOA-147Klac identified from different organisms.

**Table 5.** DNA sequences of KlacRS variants used in this study.

**Table 6.** Sequences of PCR Primers used in this study.

**Table 7.** Summary of the ALDOA-WT- and ALDOA-147Klac-interacting proteins identified from HEK293T cells.
